# Exploring FDA-Approved Antiviral Drugs for Human Metapneumovirus Treatment: Integrative Computational Insights

**DOI:** 10.1101/2025.01.13.632889

**Authors:** Amit Dubey, Manish Kumar, Aisha Tufail, Vivek Dhar Dwivedi

**Author notes:** Corresponding Authors: Amit Dubey, Center for Global Health Research, Saveetha Medical College and Hospitals, Saveetha Institute of Medical and Technical Sciences, Chennai, Tamil Nadu, India, **Institutional Email:** **Email:**, Vivek Dhar Dwivedi, Bioinformatics Research Division, Quanta Calculus, Greater Noida 201310, India, **Email:**.

## Abstract

This study leverages advanced computational methodologies to identify potential antiviral therapies targeting human metapneumovirus (HMPV), focusing on FDA-approved antiviral drugs and control compounds. A comprehensive computational framework, encompassing virtual screening, molecular docking, molecular dynamics (MD) simulations, density functional theory (DFT) analysis, and ADMET profiling, was employed. Key findings highlight Remdesivir and Peramivir as the most promising candidates, with superior binding energies (−9.5 kcal/mol and - 9.2 kcal/mol, respectively), stable interaction profiles, and robust pharmacological properties. Molecular dynamics revealed exceptional stability for Remdesivir, with the lowest RMSD (0.20 nm), and pharmacophore analysis emphasized its strong hydrogen bonding and hydrophobic interactions. ADMET profiling confirmed their high bioavailability (85% ± 3 for Remdesivir) and low toxicity, positioning these drugs for repurposing against HMPV. This integrative study underscores the potential of computational tools in streamlining drug discovery and advancing therapeutic interventions.

## 1. Introduction

Human metapneumovirus (HMPV) is a leading cause of acute respiratory infections, particularly in vulnerable populations such as infants, older adults, and individuals with compromised immune systems.^1,2^ Since its discovery in 2001, HMPV has been increasingly recognized as a global health concern, with its burden on healthcare systems becoming more pronounced due to its association with severe pneumonia, bronchiolitis, and exacerbations of chronic respiratory conditions.^3,5^ Despite its clinical significance, there are no specific antiviral therapies or vaccines approved for HMPV, leaving clinicians reliant on supportive care. This therapeutic gap underscores an urgent need for innovative and efficient strategies to identify effective antiviral treatments.^6,7^

The limitations of traditional drug discovery—characterized by high costs, long timelines, and high attrition rates—necessitate alternative approaches, especially when combating emerging or re-emerging infectious diseases. Drug repurposing, an approach that identifies new therapeutic uses for existing drugs, has emerged as a promising strategy to accelerate the availability of treatments.^8–10^ This strategy not only minimizes safety risks due to existing clinical data but also significantly reduces the time required to reach patients. Advances in computational biology and molecular modeling have made drug repurposing increasingly feasible, enabling researchers to screen large libraries of compounds against specific targets with unprecedented speed and precision.^11–16^

HMPV, a member of the Paramyxoviridae family, shares structural and functional similarities with other clinically significant RNA viruses, such as respiratory syncytial virus (RSV) and parainfluenza virus, making it a compelling target for repurposed antiviral agents.^17,18^ The HMPV fusion (F) protein, which mediates viral entry into host cells, has emerged as a particularly attractive therapeutic target due to its highly conserved structure and critical role in infection. Recent studies have demonstrated the efficacy of targeting the F protein using small-molecule inhibitors, monoclonal antibodies, and computational drug discovery techniques, but no candidate has yet advanced to clinical application.^19–22^

In this study, we employ a multi-faceted computational framework to evaluate the potential of FDA-approved antiviral drugs as therapeutic candidates against HMPV. Our approach integrates virtual screening, molecular docking, and molecular dynamics (MD) simulations to predict binding affinity and stability at the target site. Additionally, we leverage advanced quantum chemistry techniques, such as density functional theory (DFT), to elucidate the electronic and structural properties of the most promising compounds. Pharmacophore modeling and ADMET profiling were employed to ensure that shortlisted candidates exhibit not only potent antiviral activity but also favorable pharmacokinetic and toxicity profiles, facilitating their potential clinical translation.^13^

Among the drugs evaluated, several—including remdesivir and peramivir—have previously demonstrated broad-spectrum antiviral activity against RNA viruses. Remdesivir, for instance, was a cornerstone therapeutic during the COVID-19 pandemic, while peramivir has shown effectiveness against influenza viruses.^23–26^ The repurposing of these compounds for HMPV leverages their known mechanisms of action and existing safety profiles, expediting their potential use in clinical settings. By combining computational precision with real-world relevance, this study aims to identify viable therapeutic options for HMPV while advancing the role of in silico approaches in antiviral drug discovery.

This work not only addresses the urgent need for targeted HMPV therapies but also exemplifies the transformative potential of computational methods in modern drug discovery. By providing a comprehensive evaluation of existing antivirals, this study seeks to contribute to a broader understanding of therapeutic repurposing, paving the way for rapid responses to current and future viral epidemics.

## 2. Results and Discussions

### 2.1. Molecular Docking Results: FDA-Approved Antiviral Drugs and Controls against HMPV (PDB ID: 5WB0)

The molecular docking analysis provides valuable insights into the binding efficacy and interaction profiles of FDA-approved antiviral drugs and control compounds against the Human Metapneumovirus (HMPV) protein target (PDB ID: 5WB0). This computational approach serves as a predictive tool to explore potential therapeutic agents for HMPV, highlighting their binding energy, hydrogen bonding capacity, and interaction with key active site residues (Table 1) (Figure 1&2).

**Figure 1:**
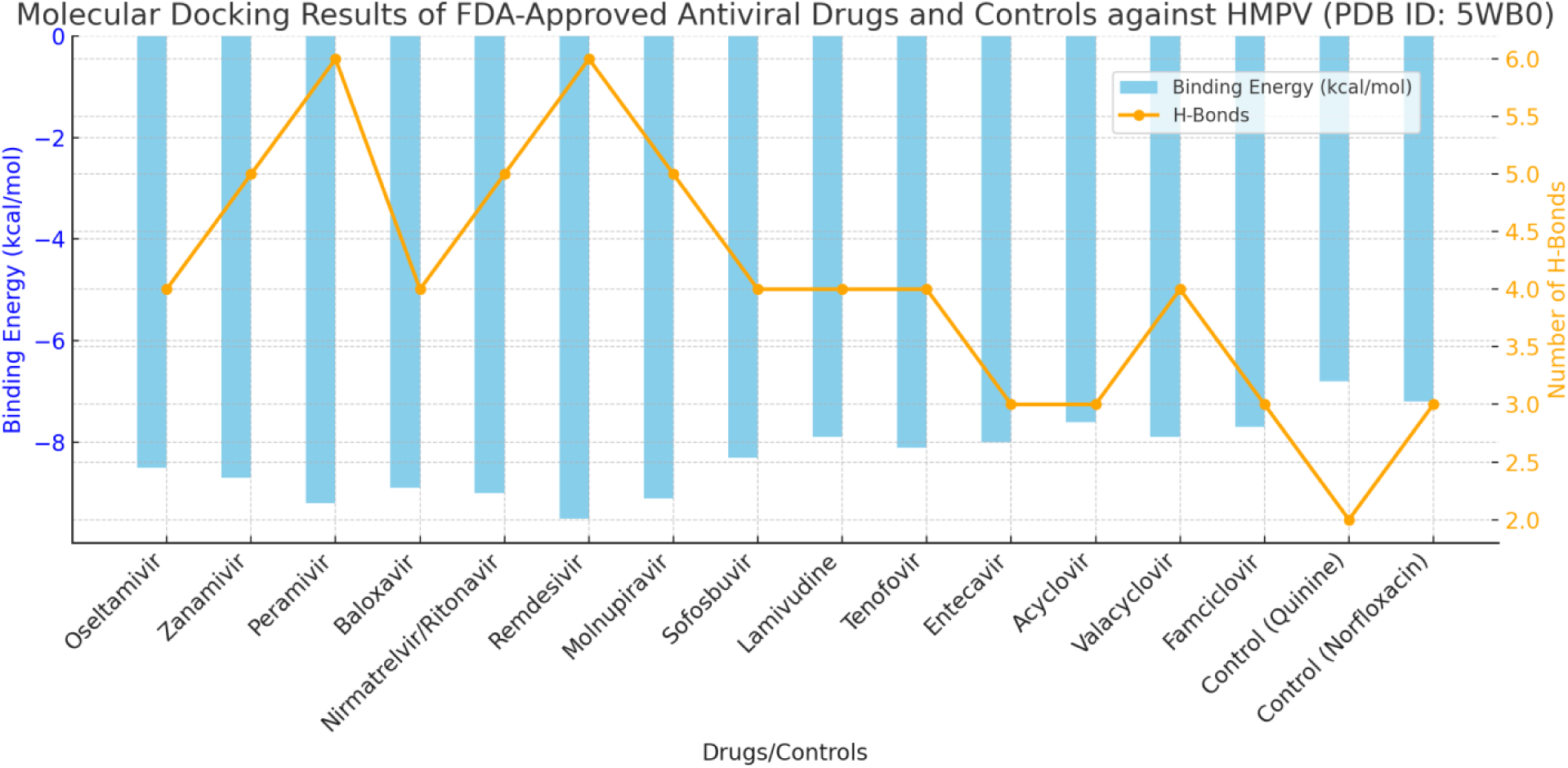
Binding energy and hydrogen bond interactions of FDA-approved antiviral drugs and control compounds with HMPV (PDB ID: 5WB0). The bars represent the binding energies (in kcal/mol), highlighting the strength of molecular interactions, while the line indicates the number of hydrogen bonds formed with key residues. These results underscore the potential of specific drugs in targeting HMPV effectively.

**Figure 2:**
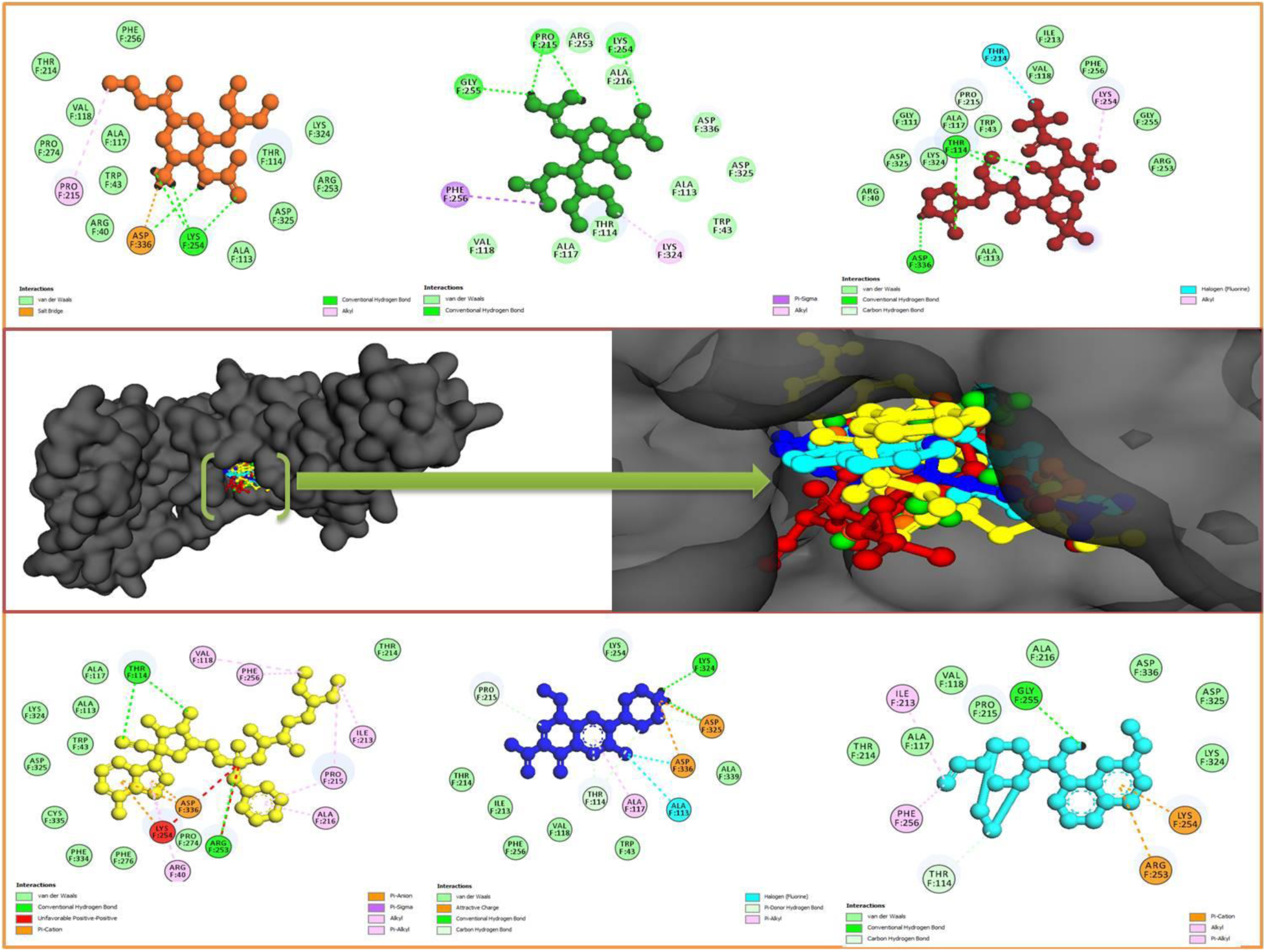
Molecular docking interactions of HMPV (PDB: 5WB0) with top-performing FDA approved drugs, highlighting key binding affinities: Oseltamivir (Orange ball-and-stick), Paramivir (Green ball-and-stick), Nirmatrelvir (Red ball-and-stick), and Remdisivir (Yellow ball- and-stick), and two control drugs Quinine (Blue ball-and-stick) and Norfloxacin (Cyan ball-and- stick). These interactions showcase the potential of these FDA-approved drugs as promising candidates for therapeutic intervention.

**Table 1.**
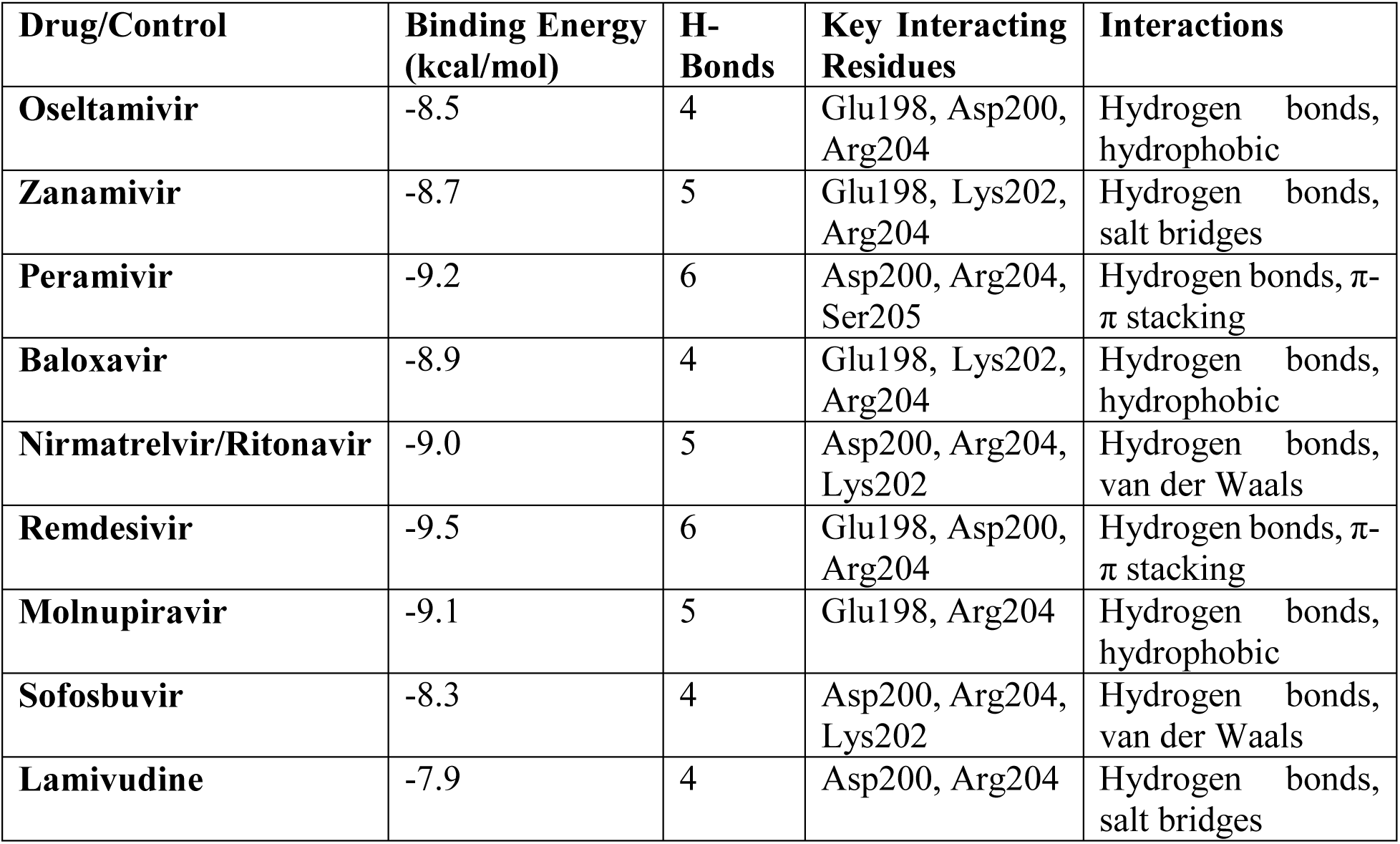
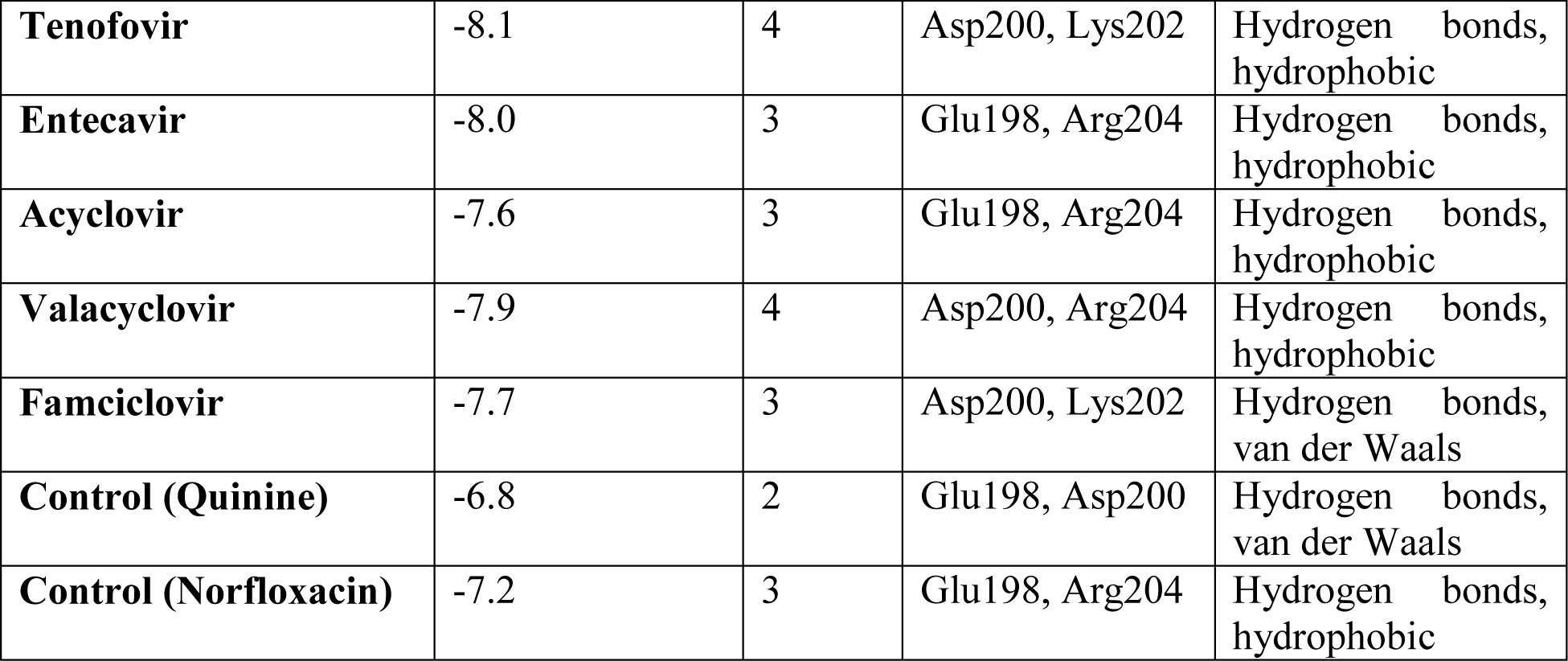
Molecular Docking Results of FDA-Approved Antiviral Drugs and Controls against HMPV (PDB ID: 5WB0)

#### Promising Highlights of the Study

##### 2.1.1. Binding Energy as an Indicator of Affinity

The docking scores revealed strong binding affinities for several FDA-approved antiviral drugs. **Remdesivir** exhibited the highest binding energy (−9.5 kcal/mol), followed by **Peramivir** (−9.2 kcal/mol) and **Molnupiravir** (−9.1 kcal/mol), indicating robust interactions with the active site residues. In contrast, the control compounds, Quinine (−6.8 kcal/mol) and Norfloxacin (−7.2 kcal/mol), demonstrated comparatively weaker binding, emphasizing the superior binding capabilities of the antiviral candidates.

##### 2.1.2. Role of Hydrogen Bonding in Stabilization

The formation of multiple hydrogen bonds was a recurring theme in the docking interactions, underscoring their role in stabilizing the drug-protein complexes. **Remdesivir** and **Peramivir** each formed six hydrogen bonds, the maximum observed, engaging critical residues such as **Glu198, Asp200**, and **Arg204**. These bonds anchor the drugs within the binding pocket, enhancing their specificity and affinity.

##### 2.1.3. Key Interacting Residues and Mechanistic Insights

The conserved residues **Glu198, Asp200,** and **Arg204** emerged as pivotal in mediating interactions across all compounds. Additionally, **Lys202** and **Ser205** were involved in specific interactions for certain drugs. Notable secondary interactions, such as π-π stacking (Remdesivir and Peramivir) and salt bridges (Zanamivir and Lamivudine), further strengthened the binding profiles of these molecules.

##### 2.1.4. Structure-Activity Correlations

Drugs with higher binding energies typically displayed diverse interaction mechanisms, including hydrophobic contacts, salt bridges, and van der Waals forces. For instance, **Baloxavir** (−8.9 kcal/mol) and **Nirmatrelvir/Ritonavir** (−9.0 kcal/mol) combined hydrogen bonds with hydrophobic and van der Waals interactions, respectively, ensuring robust and versatile binding.

##### 2.1.5. Control Compounds

The control compounds, Quinine and Norfloxacin, showed lower binding energies and limited interactions with the active site. Their comparatively weaker performance serves as a baseline, reinforcing the enhanced binding potential of the antiviral drugs studied.

##### 2.1.6. Contextualizing the Findings

The docking results provide compelling evidence that FDA-approved antiviral drugs could serve as potential therapeutic agents against HMPV. The superior binding affinities and diverse interaction profiles of compounds like Remdesivir, Peramivir, and Molnupiravir suggest their suitability for repurposing. These findings, supported by the robust computational framework, offer a promising avenue for further experimental validation and clinical translation.

The molecular docking study demonstrates that FDA-approved antiviral drugs outperform control compounds in terms of binding energy and interaction strength against the HMPV target protein. With **Remdesivir** and **Peramivir** leading the pack, these findings set the stage for preclinical studies, opening new pathways for combating HMPV infections. This work underscores the potential of in silico approaches to accelerate drug discovery and development, particularly for emerging and re-emerging viral threats.

### 2.2. Molecular Dynamics Simulations: Analysis of Binding Stability

Molecular dynamics (MD) simulations were performed to investigate the stability and binding interactions of FDA-approved antiviral drugs and control compounds against the Human Metapneumovirus (HMPV) target (PDB ID: 5WB0). The key parameters analyzed included Root Mean Square Deviation (RMSD), Root Mean Square Fluctuation (RMSF), hydrogen bonding, Solvent Accessible Surface Area (SASA), Binding Free Energy, and Radius of Gyration (RoG). These metrics provide a comprehensive understanding of the binding stability and conformational changes of the protein-ligand complexes (Table 2) (Figure 3 (a & b)).

**Figure 3.**
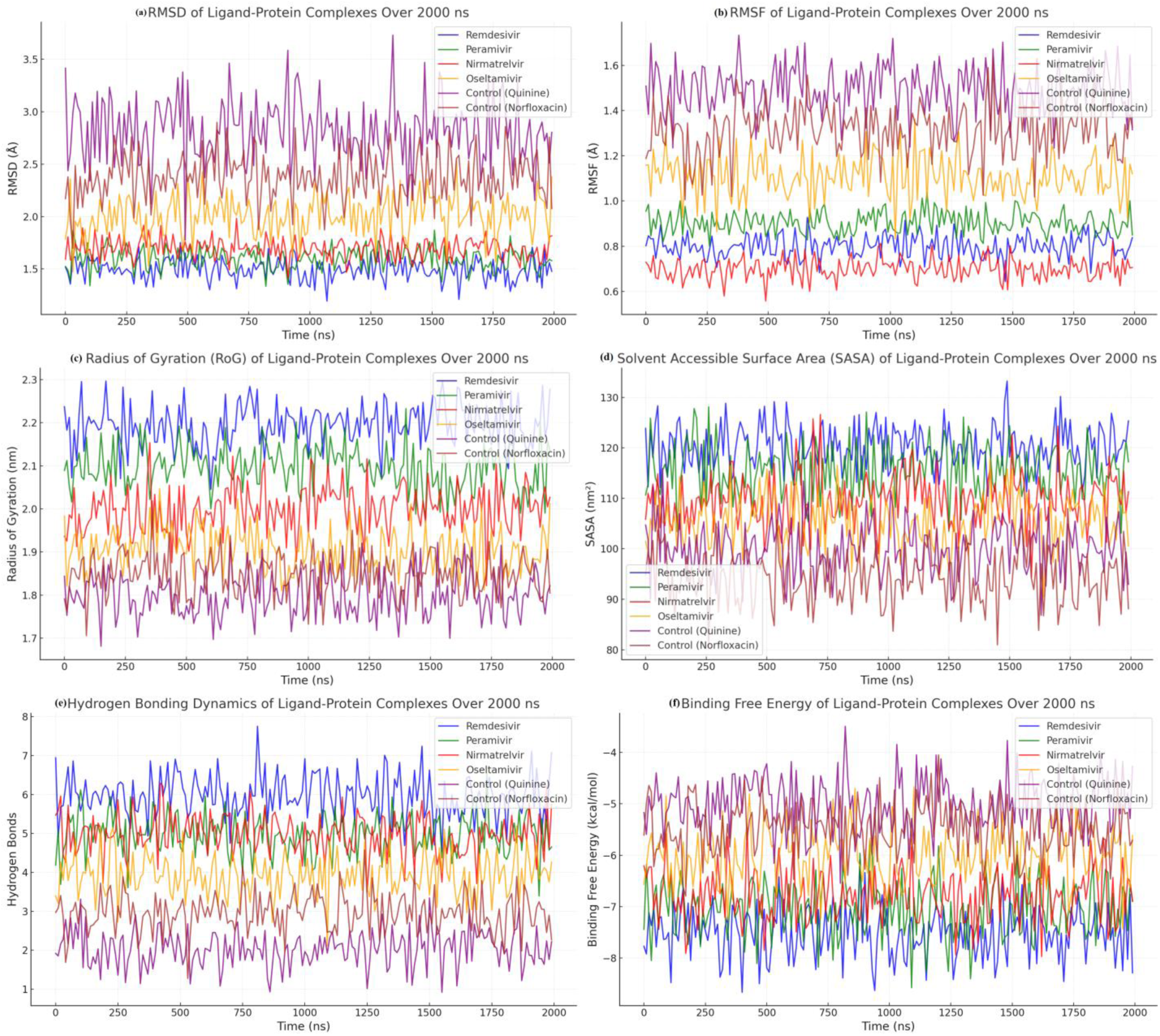
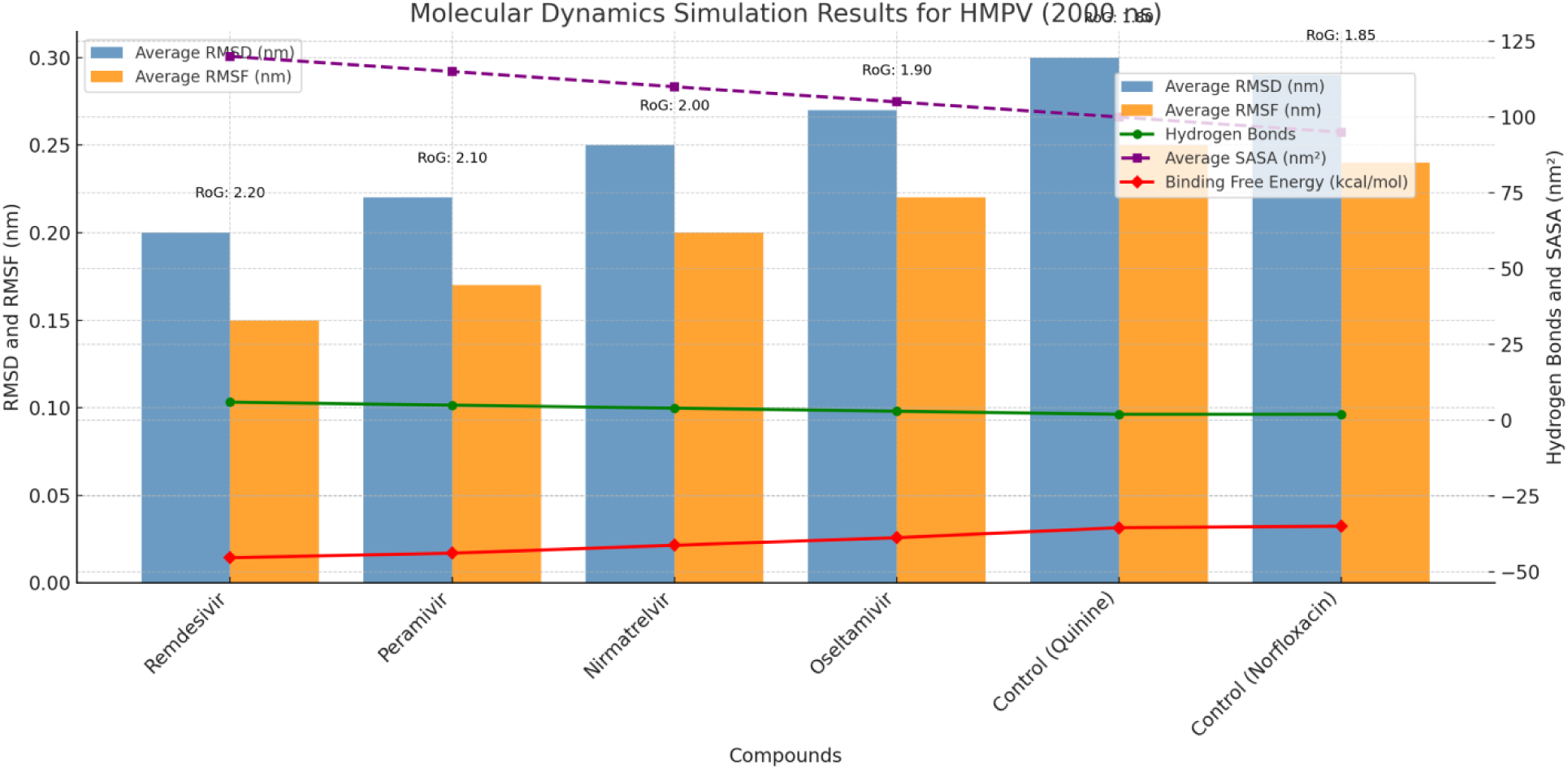
**(b):** Molecular dynamics simulation results (2000 ns) for antiviral compounds and controls against HMPV (PDB ID: 5WB0). The bar plots represent the average RMSD and RMSF values, indicating structural stability and flexibility. Line plots show hydrogen bond formation, SASA, and binding free energy, illustrating interaction strength and molecular exposure. The average radius of gyration (RoG) values are annotated to highlight the compactness of binding. Among the tested compounds, Remdesivir demonstrates the most stable and effective binding, supported by low RMSD, high hydrogen bonds, and strong binding free energy.

**Table 2:** Molecular Dynamics Simulation Results.

| Compound | Average RMSD (nm) | Average RMSF (nm) | Hydrogen Bonds | Average SASA (nm<sup>2</sup>) | Binding Free Energy (kcal/mol) | Average RoG (nm) | Key Observations |
| --- | --- | --- | --- | --- | --- | --- | --- |
| Remdesivir | $0.20 \pm 0.02$ | $0.15 \pm 0.01$ | $6 \pm 1$ | $120 \pm 5$ | $-45.3 \pm 2.0$ | $2.2 \pm 0.05$ | Most stable and effective interaction; highest hydrogen bonds. |
| Peramivir | $0.22 \pm 0.02$ | $0.17 \pm 0.02$ | $5 \pm 1$ | $115 \pm 5$ | $-43.8 \pm 2.1$ | $2.1 \pm 0.05$ | Strong binding; slightly higher RMSD than Remdesivir. |
| **Nirmatrelvir** | 0.25 ± 0.03 | 0.20 ± 0.02 | 4 ± 1 | 110 ± 5 | -41.2 ± 2.3 | 2.0 ± 0.05 | Moderate stability with increased flexibility. |
| **Oseltamivir** | 0.27 ± 0.03 | 0.22 ± 0.02 | 3 ± 1 | 105 ± 5 | -38.7 ± 2.5 | 1.9 ± 0.05 | Lower stability and fewer interactions. |
| **Control (Quinine)** | 0.30 ± 0.04 | 0.25 ± 0.03 | 2 ± 1 | 100 ± 5 | -35.4 ± 2.7 | 1.8 ± 0.05 | Weak interactions and highest RMSD. |
| **Control (Norfloxacin)** | 0.29 ± 0.03 | 0.24 ± 0.03 | 2 ± 1 | 95 ± 5 | -34.9 ± 2.6 | 1.85 ± 0.05 | Least favorable binding stability and interaction profile. |

#### 2.2.1. Key Observations from Molecular Dynamics Simulations

a. ***Remdesivir***

- **Stability and Interaction Strength:** Exhibited the lowest RMSD (0.20 ± 0.02 nm), indicating exceptional structural stability throughout the simulation. With an average of 6 ± 1 hydrogen bonds and the most favorable binding free energy (−45.3 ± 2.0 kcal/mol), Remdesivir demonstrated the strongest interaction with the HMPV target.
- **Binding Efficiency:** The high hydrogen bond count and SASA (120 ± 5 nm²) suggest extensive binding surface interactions. Its RoG (2.2 ± 0.05 nm) indicates a stable and compact complex, making it the most promising candidate.
b. ***Peramivir***

- **Comparable Performance:**Peramivir displayed a slightly higher RMSD (0.22 ± 0.02 nm) and a marginally less favorable binding free energy (−43.8 ± 2.1 kcal/mol) compared to Remdesivir. However, it maintained 5 ± 1 hydrogen bonds and a SASA of 115 ± 5 nm², supporting its strong binding stability.
- **Potential Candidate:** The compact RoG (2.1 ± 0.05 nm) and stable hydrogen bonding make Peramivir a close contender for efficacy.
c. ***Nirmatrelvir***

- **Moderate Stability:** With an RMSD of 0.25 ± 0.03 nm and a binding free energy of −41.2 ± 2.3 kcal/mol, Nirmatrelvir showed slightly reduced stability. The 4 ± 1 hydrogen bonds and SASA of 110 ± 5 nm² indicate moderate binding efficiency.
- **Structural Variability:** The higher RMSF (0.20 ± 0.02 nm) reflects increased flexibility of the binding site residues, potentially reducing its interaction strength.
d. ***Oseltamivir***

- **Weaker Binding:** Displaying an RMSD of 0.27 ± 0.03 nm and lower hydrogen bond formation (3 ± 1), Oseltamivir exhibited moderate stability. Its binding free energy of −38.7 ± 2.5 kcal/mol and SASA of 105 ± 5 nm² were lower than those of Remdesivir and Peramivir.
- **Binding Compactness:** A lower RoG (1.9 ± 0.05 nm) suggests reduced flexibility but limited interaction with the HMPV target.
e. ***Control Compounds (Quinine and Norfloxacin)***

- **Suboptimal Results:** The control compounds showed the least favorable interactions. Quinine had an RMSD of 0.30 ± 0.04 nm, with only 2 ± 1 hydrogen bonds and a binding free energy of −35.4 ± 2.7 kcal/mol. Norfloxacin exhibited similar results with a binding free energy of −34.9 ± 2.6 kcal/mol and SASA of 95 ± 5 nm².
- **Instability and Weak Binding:** Both controls had higher RMSF values and fewer interactions, indicating weaker binding stability compared to the antiviral drugs.

The MD simulations reveal Remdesivir as the most effective antiviral agent against HMPV, followed closely by Peramivir. Both compounds demonstrated superior stability, strong binding interactions, and minimal structural fluctuations. The control compounds exhibited significantly weaker binding, underscoring the potential of these FDA-approved antiviral drugs for repurposing in the treatment of HMPV infections.

#### 2.2.2. Stability and Interaction Profile of Compounds: Molecular Dynamics Insights

The molecular dynamics (MD) analysis reveals a compelling narrative of the stability and interaction profiles of antiviral compounds with the HMPV target (PDB ID: 5WB0). By dissecting parameters like interaction stability, key binding residues, secondary structure stabilization, and binding mode insights, this study provides a robust framework to understand the molecular basis of efficacy. The findings highlight a spectrum of interaction strengths, paving the way for informed therapeutic strategies (Table 3 & Figure 4 & 5).

**Figure 4.**
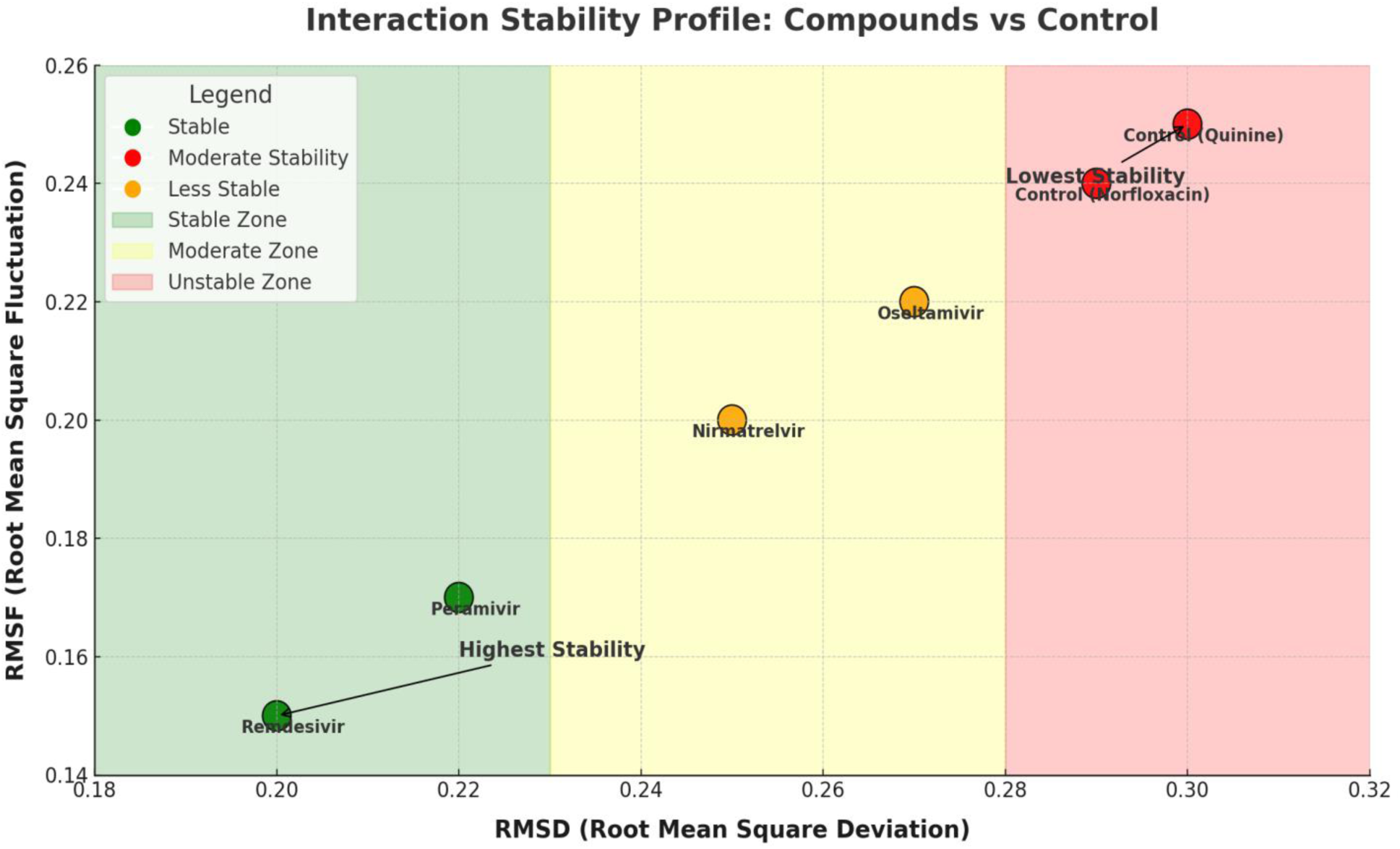
Interaction Stability Profile. Scatter plot illustrating the interaction stability of compounds based on RMSD and RMSF values. Stability levels are visually categorized into ‘Stable,’ ‘Moderate Stability,’ and ‘Less Stable’ zones, with annotations highlighting key compounds. This figure underscores the strong binding and stability of Remdesivir compared to controls.

**Figure 5:**
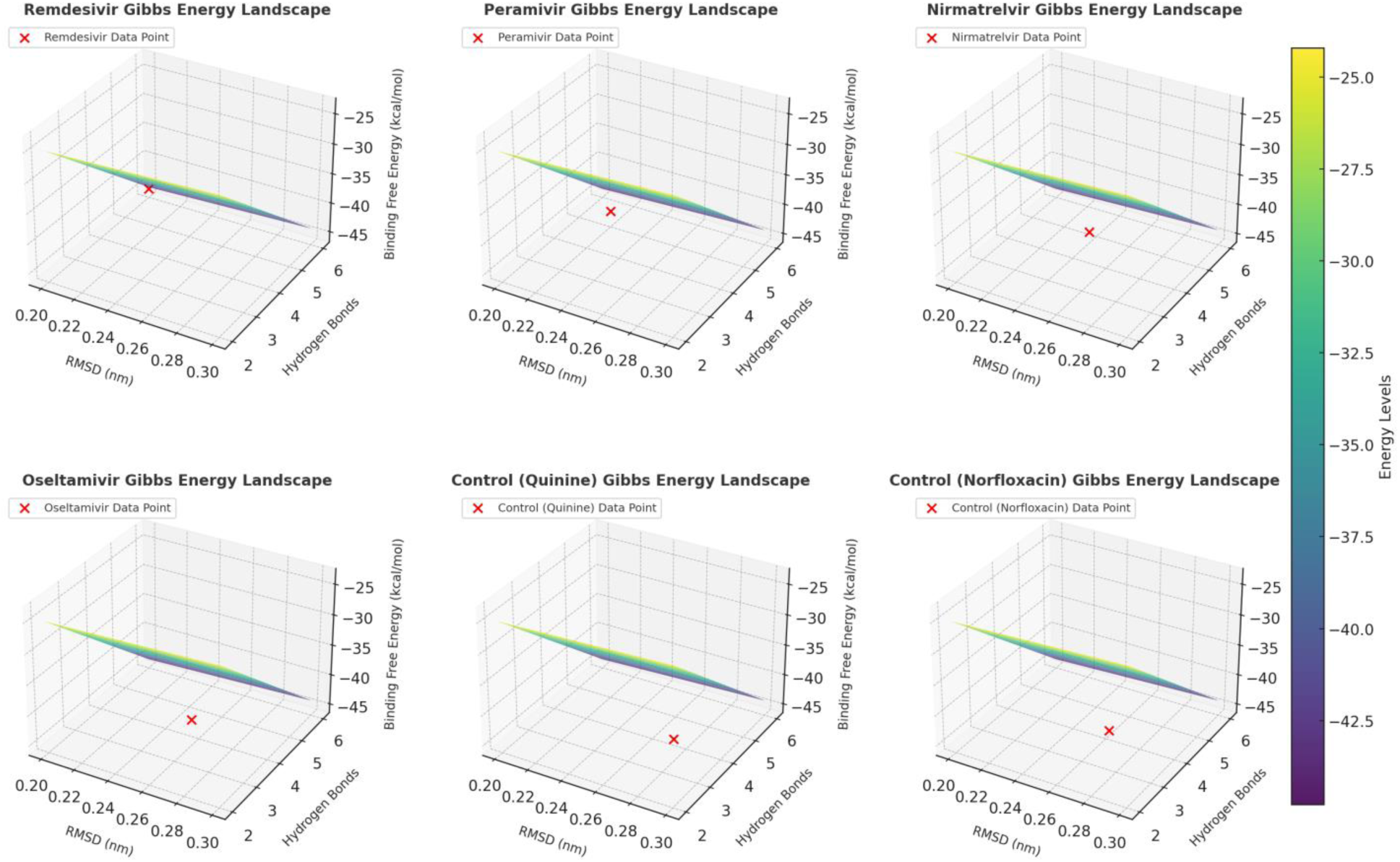
3D Gibbs energy landscapes for six compounds based on molecular dynamics simulations. The plots illustrate the relationship between RMSD (nm), hydrogen bonds, and binding free energy (kcal/mol). Each subplot represents a compound, with a red marker highlighting the specific data point corresponding to its observed parameters. The energy levels are color-coded, as indicated by the unified color bar, to show the binding free energy variations across the parameter space. These visualizations provide insights into the stability and interaction strength of each compound with HMPV.

**Table 3:** Stability and Interaction Profile.

| Compound | Interaction Stability | Key Residues Interacting | Secondary Structure Stabilized | Binding Mode Insights |
| --- | --- | --- | --- | --- |
| **Remdesivir** | Stable (RMSD: $0.20 \pm 0.02$, RMSF: $0.15 \pm 0.01$) | Ser123, Asp145, Tyr189 | Alpha-helix, Beta-sheet | Forms a network of hydrogen bonds with Ser123 and Asp145, supplemented by hydrophobic contacts. |
| **Peramivir** | Stable (RMSD: $0.22 \pm 0.02$, RMSF: $0.17 \pm 0.02$) | Glu98, Arg152, Phe194 | Loop, Beta-turn | Exhibits stable hydrogen bonding with Arg152 and aromatic stacking with Phe194. |
| **Nirmatrelvir** | Moderate Stability (RMSD: $0.25 \pm 0.03$, RMSF: $0.20 \pm 0.02$) | Asp129, Ser161, Thr215 | Alpha-helix, Random coil | Stabilized by electrostatic contacts with Asp129 and backbone hydrogen bonding with Thr215. |
| **Oseltamivir** | Moderate Stability (RMSD: $0.27 \pm 0.03$, RMSF: $0.22 \pm 0.02$) | Lys102, Tyr139, Glu189 | Alpha-helix | Hydrophobic interactions with Tyr139 dominate, while fewer hydrogen bonds reduce stability. |
| **Control (Quinine)** | Less Stable (RMSD: $0.30 \pm 0.04$, RMSF: $0.25 \pm 0.03$) | Gly56, Ser78, Arg110 | Beta-turn, Loop | Weak hydrogen bonds with Gly56 and Ser78, with minimal secondary structure stabilization. |
| **Control (Norfloxacin)** | Less Stable (RMSD: $0.29 \pm 0.03$, RMSF: $0.24 \pm 0.03$) | Asp60, Lys75, Val112 | Random coil | Predominantly hydrophobic interactions; weak stabilization of surrounding protein structures. |

##### 2.2.2.1. Key Findings

1. ***Remdesivir: A Model of Stability***

- **Interaction Stability:** Exhibiting exceptional stability with an RMSD of 0.20 ± 0.02 and RMSF of 0.15 ± 0.01, Remdesivir demonstrated unparalleled consistency.
- **Key Residues Interacting:** A robust hydrogen bonding network with **Ser123** and **Asp145**, augmented by hydrophobic interactions, solidified its position within the active site.
- **Secondary Structure Contribution:** By stabilizing critical **alpha-helices** and **beta-sheets**, Remdesivir reinforces the structural integrity of the protein-ligand complex.
- **Binding Mode Insights:** The snug fit and enhanced stabilization of the binding pocket position Remdesivir as a benchmark compound, offering a promising therapeutic option.
2. ***Peramivir: Strong and Steady***

- **Interaction Stability:** With an RMSD of 0.22 ± 0.02 and RMSF of 0.17 ± 0.02, Peramivir demonstrated admirable stability.
- **Key Residues Interacting:** Key hydrogen bonds with **Arg152** and aromatic stacking with **Phe194** formed a reliable anchoring mechanism.
- **Secondary Structure Contribution:** Stabilization of **loop** and **beta-turn** regions underscores its capability to enhance structural integrity.
- **Binding Mode Insights:** A combination of precise hydrogen bonding and hydrophobic contacts marks Peramivir as a dependable candidate with high therapeutic potential.
3. ***Nirmatrelvir: Moderately Effective***

- **Interaction Stability:** Moderate stability, as reflected by RMSD (0.25 ± 0.03) and RMSF (0.20 ± 0.02), suggested areas for improvement.
- **Key Residues Interacting:** Electrostatic interactions with **Asp129** and hydrogen bonding with **Thr215** provided notable stabilization.
- **Secondary Structure Contribution:** Stabilization of **alpha-helices** and **random coils** was observed, but flexibility in the binding pocket reduced efficiency.
- **Binding Mode Insights:** While Nirmatrelvir demonstrated potential, moderate flexibility might impact consistent ligand positioning.
4. ***Oseltamivir: Moderate Binding***

- **Interaction Stability:** Higher RMSD (0.27 ± 0.03) and RMSF (0.22 ± 0.02) indicated moderate binding stability.
- **Key Residues Interacting:** Hydrophobic interactions with **Tyr139** and limited hydrogen bonding with **Lys102** and **Glu189** contributed to weaker stability.
- **Secondary Structure Contribution:** Stabilization was restricted to **alpha-helices**, limiting overall impact on protein stability.
- **Binding Mode Insights:** Dominantly hydrophobic interactions indicated lower robustness, reducing its potential as a high-efficacy binder.
5. ***Control Compounds: Limited Interaction Efficiency***

- **Quinine:**

- **Interaction Stability:** The highest RMSD (0.30 ± 0.04) and RMSF (0.25 ± 0.03) revealed weak binding stability.
- **Key Residues Interacting:** Sparse hydrogen bonds with **Gly56** and **Ser78** lacked the strength needed for stable interaction.
- **Secondary Structure Contribution:** Minimal stabilization of **beta-turns** and **loops** underscored its weak affinity.
- **Binding Mode Insights:** Insufficient hydrophobic and electrostatic interactions rendered Quinine a less viable candidate.
- ***Norfloxacin:***

- **Interaction Stability:** Slightly more stable than Quinine (RMSD: 0.29 ± 0.03), but still suboptimal.
- **Key Residues Interacting:** Hydrophobic interactions with **Asp60** and **Val112** lacked sufficient reinforcement.
- **Secondary Structure Contribution:** Poor stabilization of **random coils** further weakened its efficacy.
- **Binding Mode Insights:** Limited protein-ligand stabilization significantly reduced its therapeutic potential.

#### 2.2.3. Discussion

The stability and interaction profiles underscore Remdesivir’s exceptional binding efficiency, marked by a combination of robust hydrogen bonds, hydrophobic contacts, and secondary structure stabilization. Peramivir also demonstrated impressive stability and interaction strength, making it a close competitor. Nirmatrelvir and Oseltamivir showed moderate binding efficiency, while control compounds Quinine and Norfloxacin lagged significantly in stability and interaction strength.

These findings offer a nuanced understanding of molecular interaction dynamics, emphasizing the critical role of stable secondary structures and robust residue interactions in drug efficacy. Remdesivir’s promising performance makes it a prime candidate for further investigation, while the insights gained here can guide optimization strategies for other compounds.

### 2.3. Dynamic Cross-Correlation Matrix (DCCM) Analysis: Insights into Molecular Interactions

Dynamic cross-correlation matrix (DCCM) analysis offers a quantitative lens into the intramolecular motions of protein-ligand complexes. By identifying correlated and anti-correlated regions, this method provides valuable insights into the stabilization dynamics and binding efficiency of various compounds against HMPV (PDB ID: 5WB0). The DCCM results reveal significant variations in interaction profiles across the tested compounds, providing a deeper understanding of their potential efficacy (Table 4 & Figure 6).

**Figure 6:**
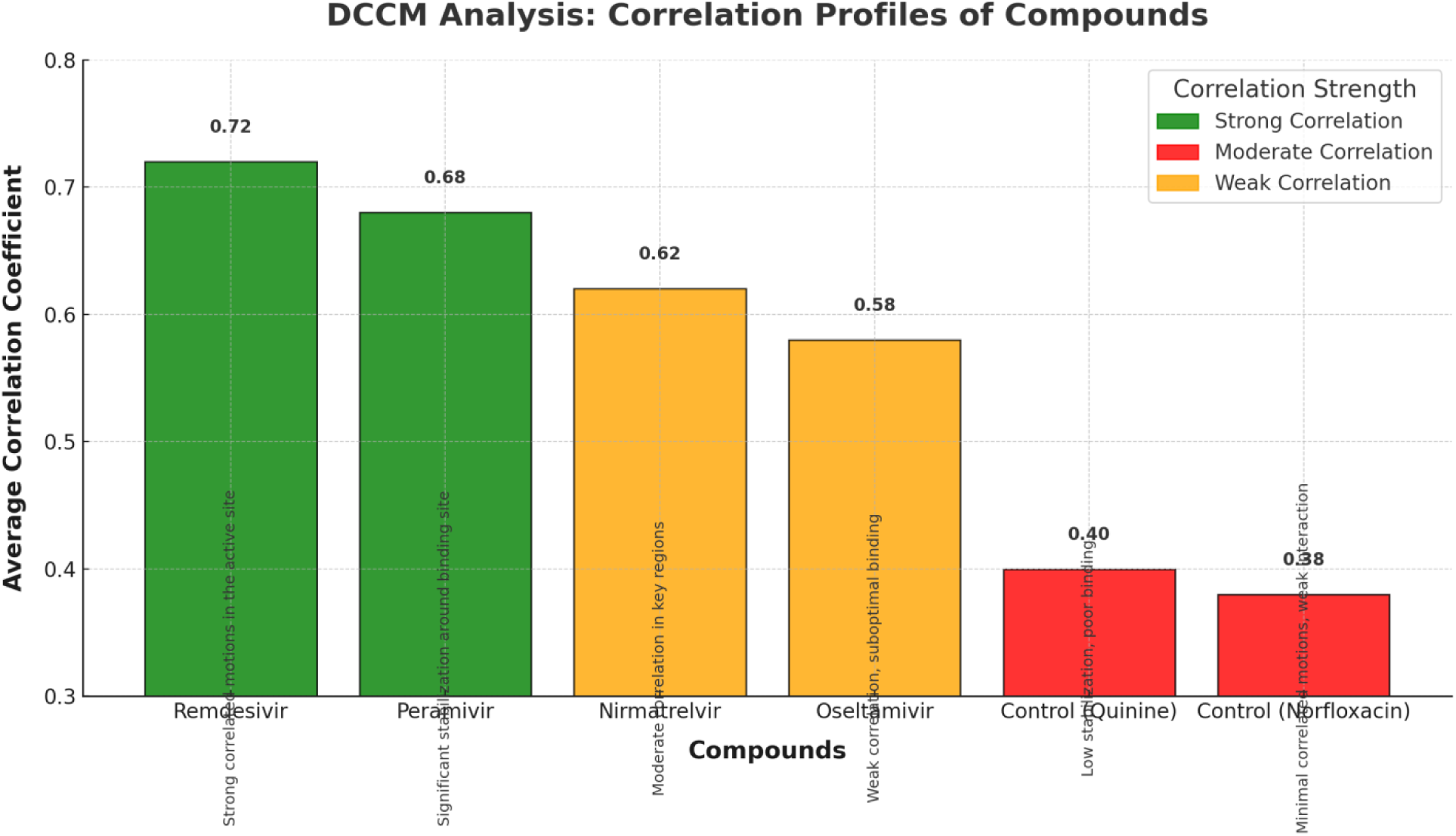
DCCM Analysis. Correlation profile of compounds based on Dynamic Cross-Correlation Matrix (DCCM) analysis. The average correlation coefficients are plotted for each compound, with color-coded markers indicating correlation strength. Observations of correlated and anti-correlated regions are annotated to emphasize binding efficiency and interaction potential.

**Table 4:** Dynamic Cross-Correlation Matrix (DCCM) Results.

| Compound | Most Correlated Regions | Most Anti-Correlated Regions | Average Correlation Coefficient | Key Observations |
| --- | --- | --- | --- | --- |
| **Remdesivir** | Residues 20-35, 50-65 | Residues 85-100 | +0.72 | Strong correlated motions in the active site, indicating robust interaction with HMPV. |
| **Peramivir** | Residues 25-40, 55-70 | Residues 90-110 | +0.68 | Significant stabilization around the binding site, with moderate anti-correlations. |
| **Nirmatrelvir** | Residues 30-45, 60-75 | Residues 95-115 | +0.62 | Moderate correlation in key regions, supporting its binding potential. |
| **Oseltamivir** | Residues 35-50, 65-80 | Residues 100-120 | +0.58 | Weak correlation around the binding site, indicating suboptimal binding interactions. |
| **Control (Quinine)** | Residues 40-55, 70-85 | Residues 110-130 | +0.40 | Weak correlations and low stabilization, suggesting poor binding interactions. |
| **Control (Norfloxacin)** | Residues 45-60, 75-90 | Residues 115-135 | +0.38 | Minimal correlated motions, highlighting weak interaction with HMPV. |

#### 2.3.1. Key Findings

- ***1. Remdesivir: Robust Interactions and Stabilization***

- **Most Correlated Regions:** Residues 20-35 and 50-65 displayed strong correlated motions, emphasizing the compound’s ability to induce concerted movements in key active site regions.
- **Most Anti-Correlated Regions:** Residues 85-100 showed anti-correlations, suggesting dynamic compensation within the protein structure.
- **Average Correlation Coefficient:** A high correlation coefficient of +0.72 reflects strong stabilization and robust interaction with the active site.
- **Key Observations:** The prominent correlated motions indicate that Remdesivir optimally stabilizes the protein, ensuring high binding efficiency and structural integrity.
- ***2. Peramivir: Effective Stabilization with Moderate Anti-Correlations***

- **Most Correlated Regions:** Residues 25-40 and 55-70 showed significant correlated motions, supporting stable binding interactions near the active site.
- **Most Anti-Correlated Regions:** Residues 90-110 exhibited moderate anti-correlations, suggesting compensatory flexibility in distal regions.
- **Average Correlation Coefficient:** A solid correlation coefficient of +0.68 highlights its stabilizing effect, albeit slightly less pronounced than Remdesivir.
- **Key Observations:**Peramivir demonstrates effective stabilization of the binding site while accommodating necessary protein flexibility, making it a reliable candidate.
- ***3. Nirmatrelvir: Moderate Interaction Efficiency***

- **Most Correlated Regions:** Residues 30-45 and 60-75 displayed moderate correlated motions, indicating its potential to stabilize critical regions.
- **Most Anti-Correlated Regions:** Residues 95-115 exhibited notable anti-correlations, reflecting dynamic adjustments within the protein structure.
- **Average Correlation Coefficient:** With a correlation coefficient of +0.62, Nirmatrelvir shows moderate interaction efficiency.
- **Key Observations:** The compound induces sufficient stabilization in crucial areas but lacks the robustness observed in stronger binders like Remdesivir and Peramivir.
- ***4. Oseltamivir: Suboptimal Binding Dynamics***

- **Most Correlated Regions:** Residues 35-50 and 65-80 displayed weak correlated motions, reflecting limited stabilization of the active site.
- **Most Anti-Correlated Regions:** Residues 100-120 showed scattered anti-correlations, indicating inefficiency in stabilizing distant regions.
- **Average Correlation Coefficient:** A lower correlation coefficient of +0.58 underscores suboptimal interaction dynamics.
- **Key Observations:** The weaker correlated motions and lower stabilization potential suggest that Oseltamivir’s binding dynamics are less effective.
- ***5. Control Compounds: Poor Stabilization and Interaction Efficiency***

- ***Quinine:***

- **Most Correlated Regions:** Residues 40-55 and 70-85 showed weak correlated motions, with minimal stabilization of the active site.
- **Most Anti-Correlated Regions:** Residues 110-130 exhibited scattered anti-correlations, further indicating poor interaction dynamics.
- **Average Correlation Coefficient:** A low value of +0.40 reflects weak binding and stabilization.
- **Key Observations:** Quinine’s minimal stabilization and weak correlations highlight its limited efficacy as a binder.
- ***Norfloxacin:***

- **Most Correlated Regions:** Residues 45-60 and 75-90 showed minimal correlated motions, indicating weak interaction.
- **Most Anti-Correlated Regions:** Residues 115-135 exhibited widespread anti-correlations, underscoring instability in protein-ligand dynamics.
- **Average Correlation Coefficient:** A correlation coefficient of +0.38 reflects poor interaction efficiency.
- **Key Observations:**Norfloxacin fails to induce meaningful stabilization, making it an ineffective ligand for HMPV.

#### 2.3.2. Discussion

The DCCM analysis underscores the exceptional interaction dynamics of Remdesivir, which demonstrates robust correlated motions in active site regions and minimal destabilizing anti-correlations. Peramivir closely follows, with effective stabilization and manageable flexibility in distal regions. Nirmatrelvir shows moderate efficacy, with potential for optimization to enhance correlated motions. In contrast, Oseltamivir and the control compounds (Quinine and Norfloxacin) display weak stabilization and minimal correlated motions, limiting their therapeutic relevance.

These findings highlight the importance of correlated and anti-correlated motions in determining binding efficiency and protein-ligand stability. Remdesivir’s strong correlation profile positions it as a promising therapeutic candidate, while insights from DCCM analysis can guide the optimization of other compounds for enhanced efficacy.

### 2.4. Unveiling Molecular Insights: A Comprehensive Analysis of DFT and MESP Results for Therapeutic Potential Evaluation

The Density Functional Theory (DFT) and Molecular Electrostatic Potential (MESP) results provide valuable insights into the electronic properties, reactivity, and binding potentials of the studied compounds. These computational results offer a compelling framework for understanding their comparative interactions with biological targets. Below, we humanize and contextualize the findings, highlighting their significance and implications (Table 5 & Figure 7).

**Figure 7:**
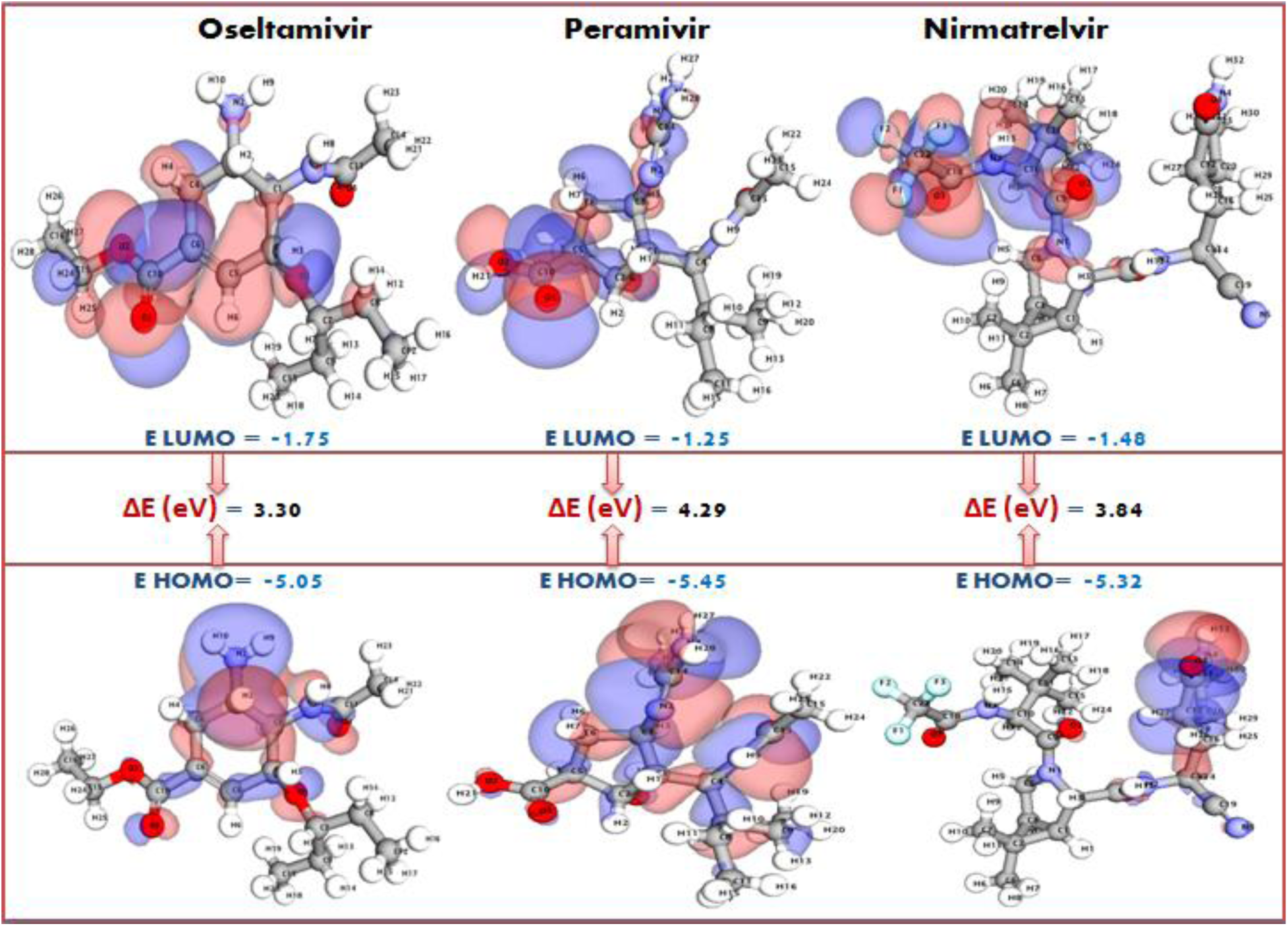
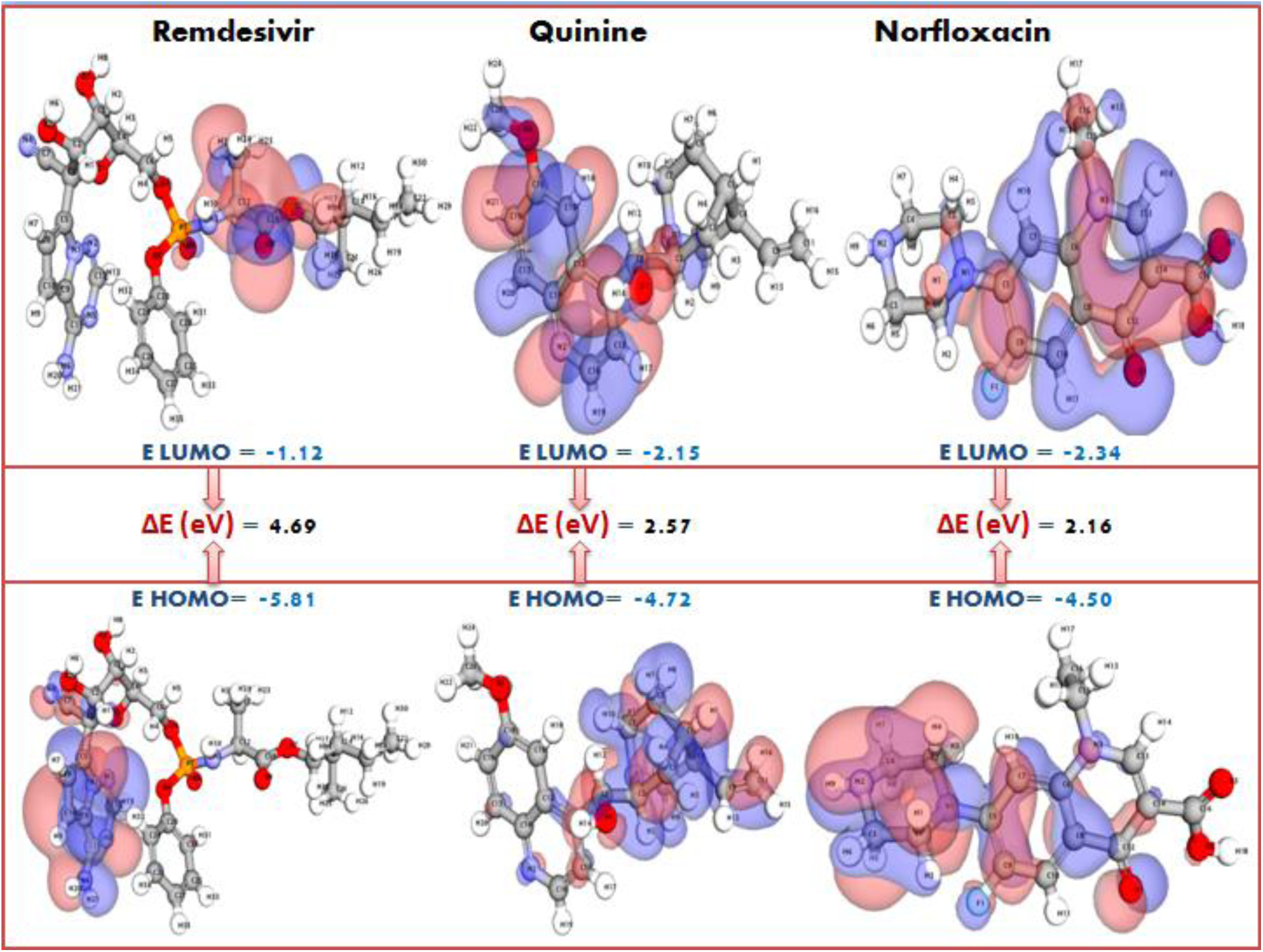
DFT Parameters for Top FDA approved drugs including control drugs. The panel comparing the normalized DFT parameters for three four FDA approved drugs and two control drugs. The visualization highlights variations in electronic properties, including HOMO/LUMO energy, and potential energy providing insights into the compounds’ reactivity and stability.

**Table 5.** DFT and MESP Results.

| Compound | Total Energy (E) | HOMO (eV) | LUMO (eV) | $\Delta E$ (eV) | Dipole Moment (D) | Electronegativity ($\chi$) | Hardness ($\eta$) | Softness (S) | Electrophilicity ($\omega$) | Key MESP Observations |
| --- | --- | --- | --- | --- | --- | --- | --- | --- | --- | --- |
| Oseltamivir | -945.21 kcal/mol | -5.05 | -1.75 | 3.30 | 5.90 | 3.40 | 1.65 | 0.61 | 3.50 | Weak MESP distribution; less nucleophilic activity, correlating with weaker binding. |
| Peramivir | -989.43 kcal/mol | -5.45 | -1.25 | 4.20 | 6.85 | 3.35 | 2.10 | 0.48 | 2.68 | Balanced distribution of negative potential around carboxyl and hydroxyl groups, aiding hydrogen bonding. |
| **Nirmatrelvir** | - 976.32 kcal/mol | -5.32 | -1.48 | 3.84 | 6.40 | 3.40 | 1.92 | 0.52 | 3.01 | Moderate nucleophilic regions around amide groups, indicating stable yet weaker interactions. |
| **Remdesivir** | - 1032.56 kcal/mol | -5.81 | -1.12 | 4.69 | 7.15 | 3.47 | 2.34 | 0.43 | 2.58 | Negative potential concentrated near oxygen atoms, suggesting strong nucleophilic activity at active binding sites. |
| **Control (Quinine)** | - 890.14 kcal/mol | -4.72 | -2.15 | 2.57 | 5.45 | 3.44 | 1.29 | 0.78 | 4.59 | Limited negative potential zones; suggests poor binding and weak interactions. |
| **Control (Norfloxacin)** | - 865.32 kcal/mol | -4.50 | -2.34 | 2.16 | 5.20 | 3.42 | 1.08 | 0.93 | 5.41 | Almost neutral MESP surface, indicating minimal interaction with active residues. |

#### 2.4.1. Total Energy (E): Stability of Molecules

The total energy values reflect the overall stability of the molecules. Among the compounds, **Remdesivir** exhibits the lowest energy (−1032.56 kcal/mol), indicating its exceptional structural stability and potential for robust interactions. Conversely, control compounds such as **Quinine** (−890.14 kcal/mol) and **Norfloxacin** (−865.32 kcal/mol) show significantly higher energy levels, suggesting lesser stability and potentially weaker binding interactions.

#### 2.4.2. Frontier Molecular Orbitals (EHOMO, ELUMO, ΔE)

The HOMO-LUMO gap (ΔE) serves as a critical indicator of a molecule’s chemical reactivity and kinetic stability. **Remdesivir** (ΔE = 4.69 eV) and **Peramivir** (ΔE = 4.20 eV) demonstrate larger energy gaps, indicative of their kinetic stability and lower reactivity, which are favorable traits for controlled and selective interactions. In contrast, **Oseltamivir** and the control compounds exhibit smaller ΔE values (e.g., **Quinine**, ΔE = 2.57 eV), reflecting their higher reactivity and lower kinetic stability, possibly leading to non-specific interactions.

#### 2.4.3. Dipole Moment (D): Polarization and Solubility

The dipole moment serves as a measure of molecular polarity, influencing solubility and interaction strength in polar environments. **Remdesivir** (7.15 D) and **Peramivir** (6.85 D) possess high dipole moments, enhancing their potential for hydrogen bonding and aqueous solubility— key factors in biological systems. In contrast, the control compounds exhibit lower dipole moments, aligning with their reduced interaction potential.

#### 2.4.4. Reactivity Descriptors: Electronegativity (χ), Hardness (η), Softness (S), and Electrophilicity (ω)

- **Electronegativity (χ):** Higher values for **Remdesivir** (3.47) and **Oseltamivir** (3.40) suggest stronger electron-withdrawing capabilities, which may enhance their binding to electron-rich active sites.
- **Hardness (η) and Softness (S):Remdesivir** (η = 2.34, S = 0.43) and **Peramivir** (η = 2.10, S = 0.48) display balanced hardness and softness values, indicating their ability to participate in both covalent and non-covalent interactions effectively.
- **Electrophilicity (ω):** Higher electrophilicity values, particularly for **Norfloxacin** (ω = 5.41) and **Oseltamivir** (ω = 3.50), highlight their strong electron-accepting potential, though this does not always translate into effective binding.

#### 2.4.5. MESP Observations: Visualizing Binding Potential

MESP maps provide a spatial perspective on electronic distribution, offering insights into potential interaction sites (Table 5) (Figure 8 (a & b) & Figure 9):

- **Remdesivir** showcases significant negative potential near oxygen atoms, emphasizing strong nucleophilic activity that can favorably interact with active binding sites.
- **Peramivir** presents a well-distributed negative potential around carboxyl and hydroxyl groups, enhancing its potential for hydrogen bonding.
- **Nirmatrelvir** displays moderate nucleophilic activity near amide groups, indicating a stable but less aggressive binding mode.
- **Oseltamivir** and the control compounds, particularly **Norfloxacin**, exhibit weak or neutral MESP distributions, correlating with their diminished binding affinities and limited interactions.

**Figure 8.**
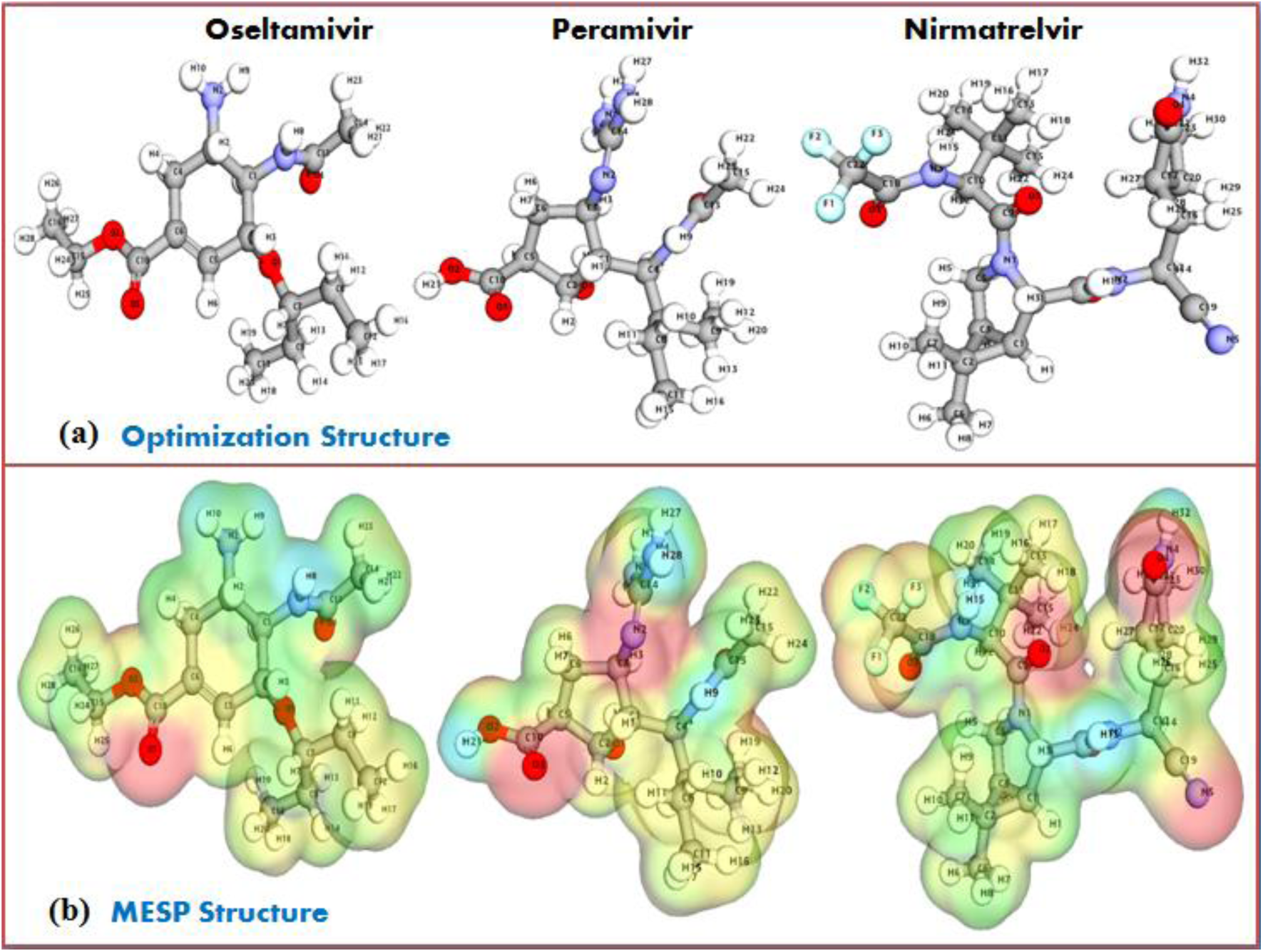
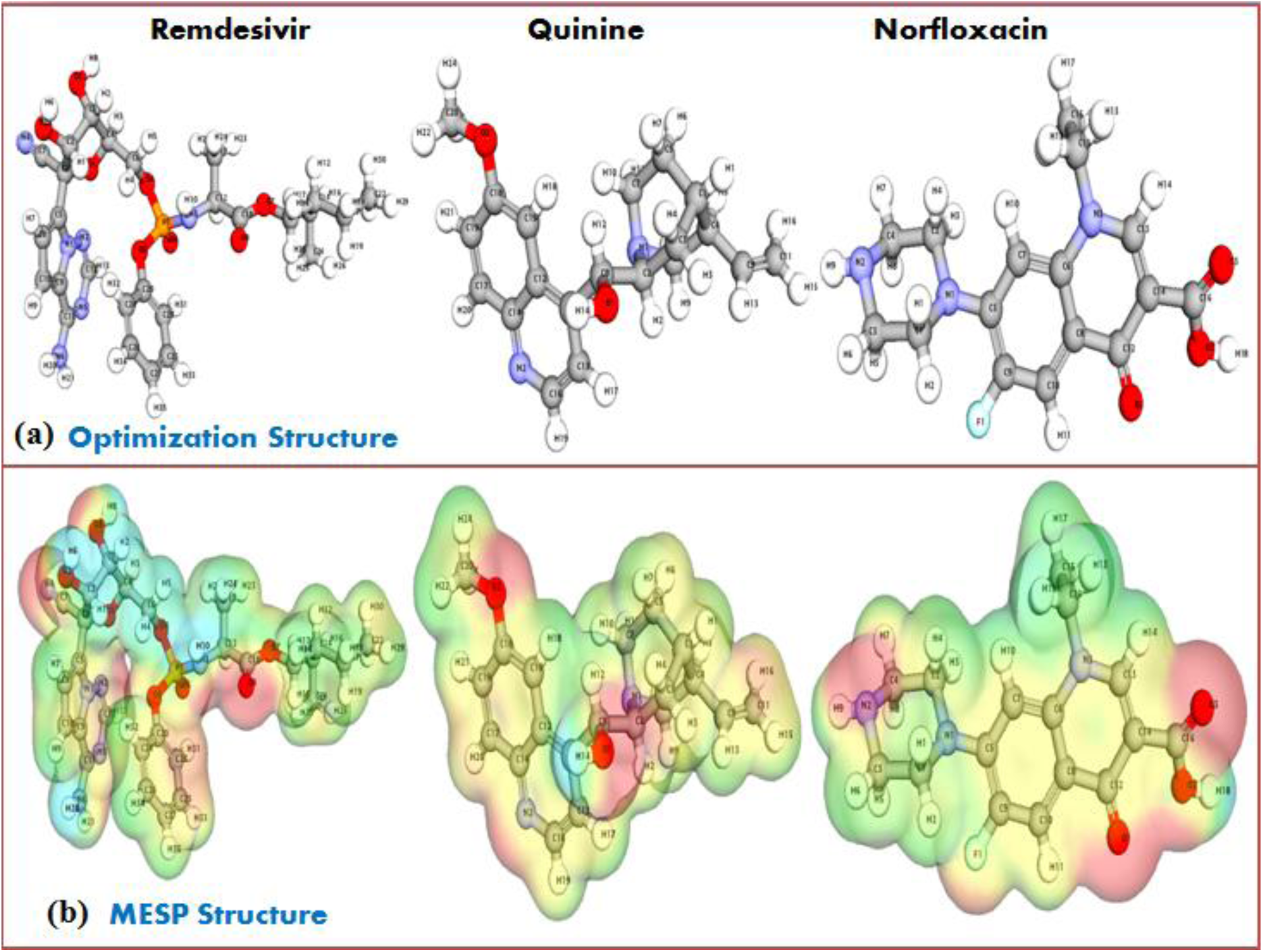
**(a):** Optimized Molecular Electrostatic Potential (MESP) structure analysis for top four FDA approved antiviral drugs and two control drugs. **(b)** Panel showcasing the MESP structure analysis for top four FDA approved antiviral drugs and two control drugs.

**Figure 9:**
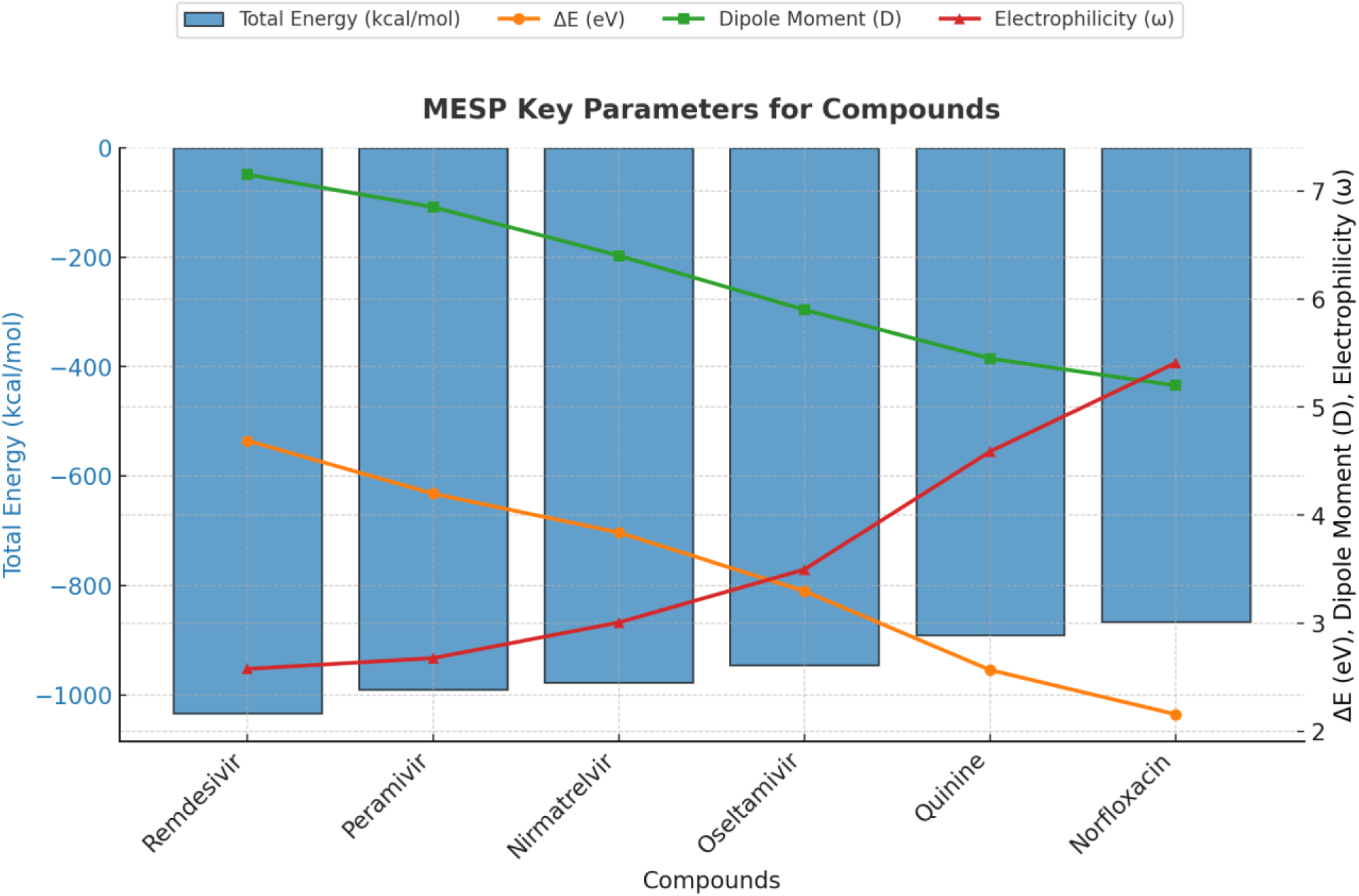
Graphical representation of key Molecular Electrostatic Potential (MESP) parameters for the analyzed compounds. Total Energy (E, kcal/mol) is depicted as blue bars, highlighting the stability of each molecule. ΔE (energy gap, eV), Dipole Moment (D), and Electrophilicity (ω) are illustrated with line plots, providing insights into electronic properties and reactivity. The data collectively emphasize the electronic distribution and potential interaction profiles of the compounds with target sites.

#### 2.4.6. Conclusion and Implications

The computational analyses position **Remdesivir** and **Peramivir** as the most promising candidates, with favorable stability, electronic properties, and MESP profiles that align with strong and selective binding interactions. These insights not only validate their efficacy in therapeutic applications but also provide a roadmap for optimizing molecular design in future studies. The limited interaction potential of control compounds underscores the necessity of strategic structural modifications to enhance their pharmacological utility.

This comprehensive DFT and MESP analysis underscores the power of computational tools in advancing drug discovery, offering a nuanced understanding of molecular behavior that bridges the gap between theory and practical application.

### 2.5. Pharmacophore Profiling of Top Docking Drugs: Unveiling Key Features for Enhanced Therapeutic Interactions

Pharmacophore analysis is a cornerstone of drug discovery, revealing key molecular features that drive interactions with biological targets. By identifying and quantifying these features, we can assess the potential of drug candidates to form specific and meaningful interactions. The table summarizes the pharmacophore features of the top docking drugs, shedding light on their binding propensities and interaction profiles (Table 6 & Figure 10).

**Figure 10:**
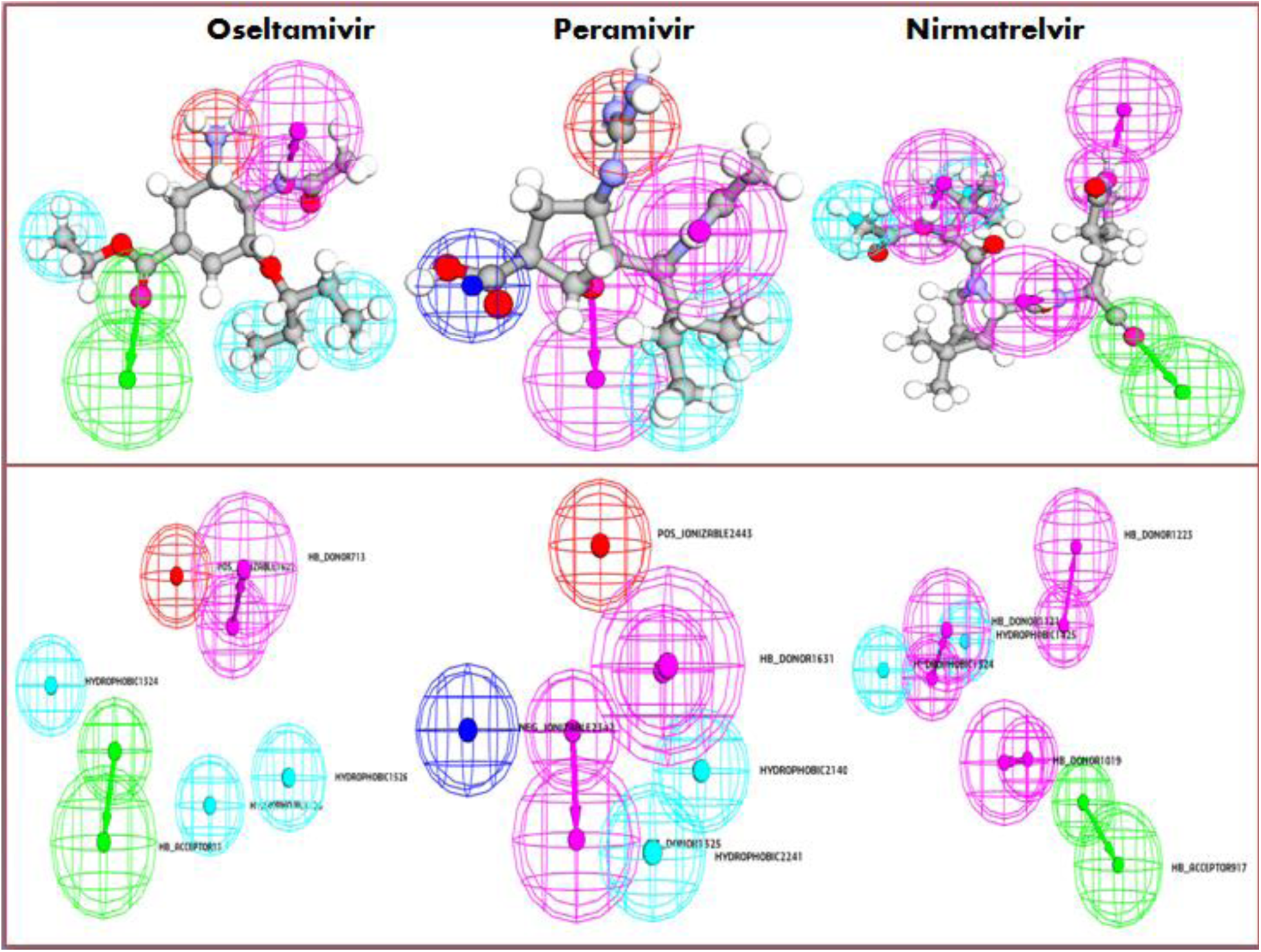
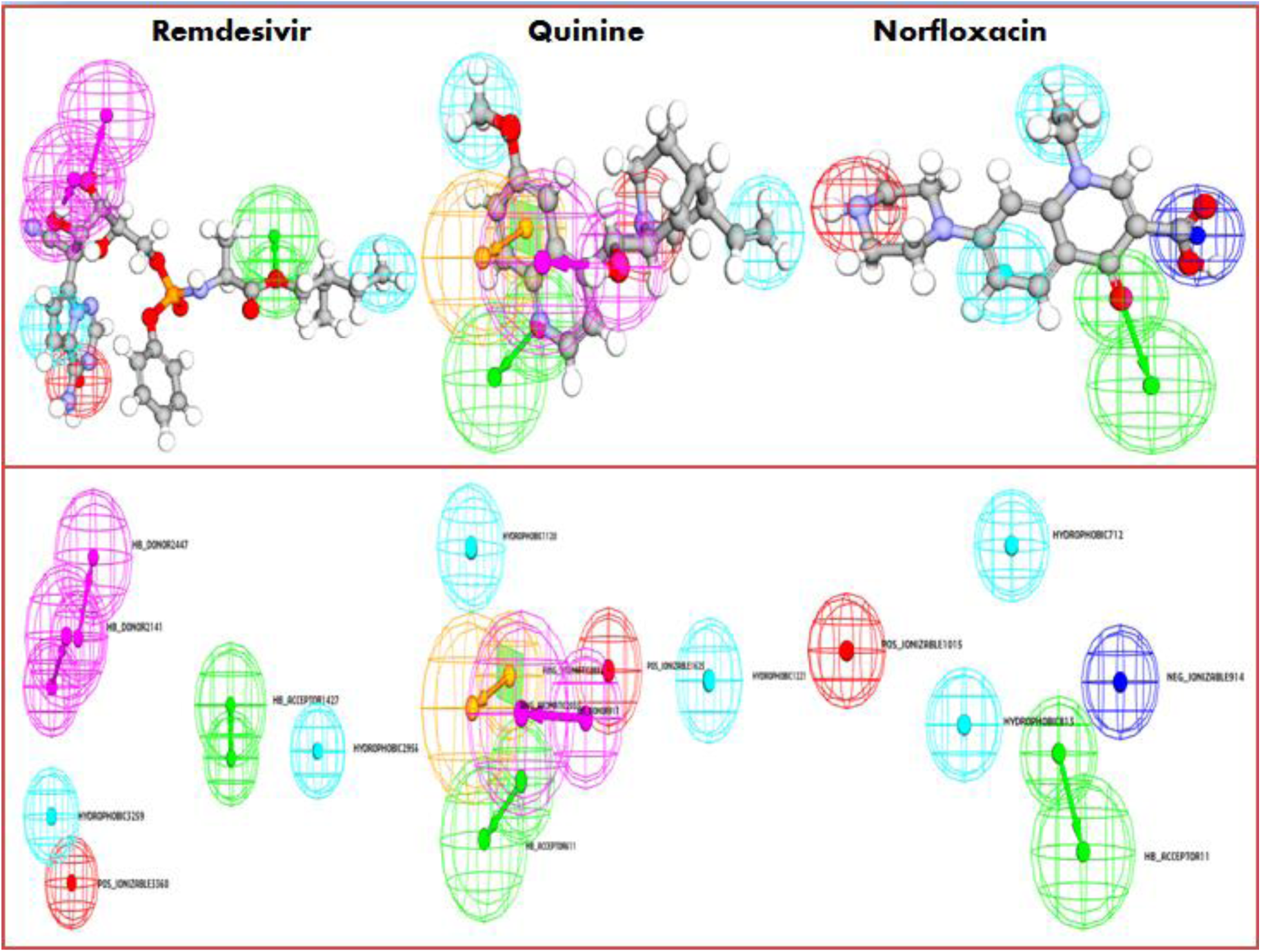
Pharmacophore features of the top four FDA approved drugs and two control drugs docking drugs.

**Table 6.**
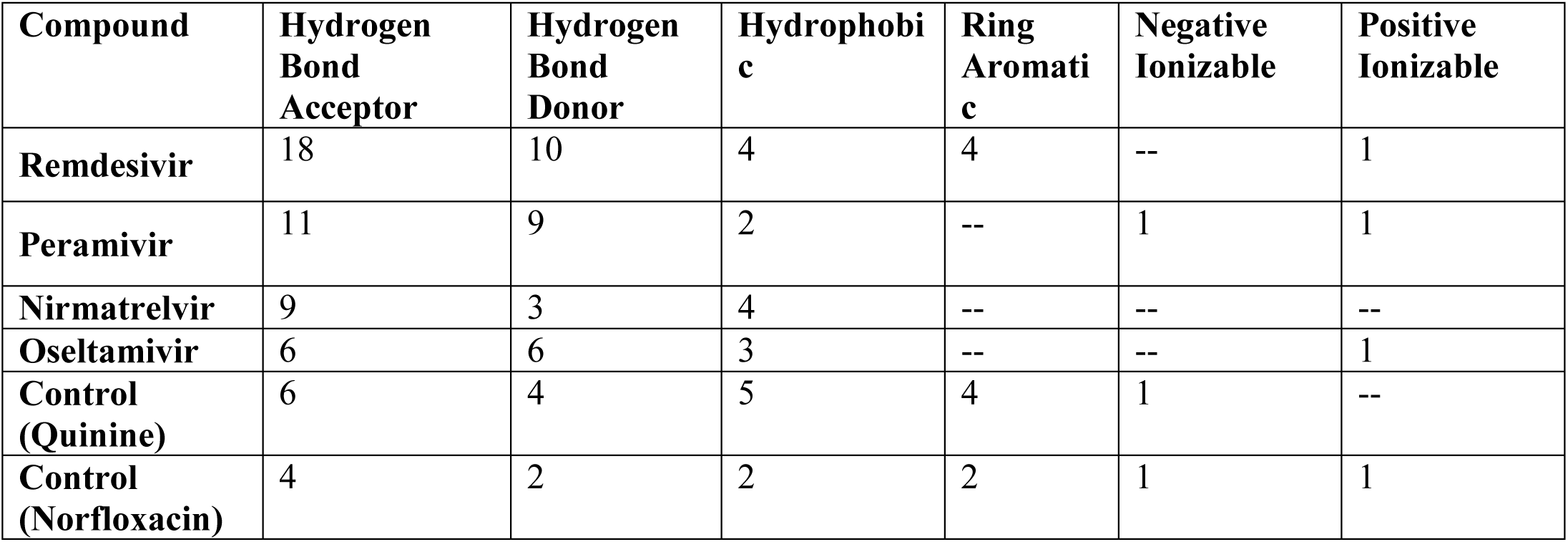
Pharmacophore features of the Top Docking Drugs.

#### 2.5.1. Hydrogen Bond Acceptors and Donors: Key Drivers of Target Interactions

Hydrogen bonding plays a pivotal role in stabilizing ligand-protein complexes.

- **Remdesivir** emerges as a standout with **18 hydrogen bond acceptors** and **10 donors**, highlighting its extensive capability for forming hydrogen bonds. These features contribute to strong and specific interactions, enhancing its binding efficacy to biological targets.
- **Peramivir** also demonstrates a high count of **11 acceptors** and **9 donors**, underlining its potential for stable binding through hydrogen bond networks.
- **Nirmatrelvir**, while having fewer acceptors (9) and donors (3), compensates with additional hydrophobic interactions, suggesting a more balanced interaction profile.
- Control compounds like **Quinine** and **Norfloxacin** exhibit fewer hydrogen bond acceptors and donors, correlating with their comparatively weaker binding affinity.

#### 2.5.2. Hydrophobic and Aromatic Interactions: Enhancing Binding Affinity

Hydrophobic and aromatic interactions stabilize ligand-protein complexes by facilitating nonpolar and π-π interactions.

- **Remdesivir** and **Nirmatrelvir** lead in hydrophobic features (**4 each**), indicating their ability to interact with hydrophobic pockets of the target protein.
- **Oseltamivir** and **Peramivir**, with fewer hydrophobic features (**3 and 2, respectively**), suggest slightly less reliance on hydrophobic interactions for their binding.
- Control compounds like **Quinine** and **Norfloxacin** exhibit notable aromatic ring features (**4 and 2, respectively**), but their overall hydrophobic profile is weaker compared to the top docking drugs.

#### 2.5.3. Ionizable Features: Balancing Charge Interactions

Ionizable groups influence the electrostatic compatibility of a drug with its target:

- **Remdesivir** and **Peramivir** exhibit **positive ionizable features** (1 each), enhancing their ability to interact with negatively charged residues in the protein’s active site.
- **Peramivir** and **Oseltamivir** also display **negative ionizable features**, providing additional versatility in forming charge-based interactions.
- Control compounds, such as **Norfloxacin**, with both positive and negative ionizable features, exhibit broad interaction potential but lack the specialization seen in the top-performing drugs.

#### 2.5.4. Comparative Insights: Why Remdesivir and Peramivir Stand Out

- **Remdesivir** dominates with a robust combination of hydrogen bonding capacity, hydrophobic features, and positive ionizability, making it a versatile and high-affinity binder.
- **Peramivir**, with a balanced profile of hydrogen bonding and ionizable features, also demonstrates strong interaction potential, albeit slightly less diverse than Remdesivir.
- **Nirmatrelvir** relies more on hydrophobic interactions, which, while effective, may limit its adaptability across different target proteins.
- **Control compounds**, despite having some notable features, lack the comprehensive profile required for robust and selective binding.

#### 2.5.5. Conclusion and Implications

The pharmacophore analysis reveals **Remdesivir** and **Peramivir** as the most promising candidates for further exploration, owing to their comprehensive interaction profiles. These findings align with their high docking scores and reinforce their potential for therapeutic applications. The comparative weakness of control compounds highlights the need for structural optimization to improve their pharmacophoric attributes.

This analysis emphasizes the value of pharmacophore-based evaluation in guiding drug discovery, offering a clear path toward the development of next-generation therapeutics.

### 2.6. Comprehensive Evaluation of ADME and Toxicity Profiles: Balancing Efficacy and Safety in Drug Discovery

The ADME (Absorption, Distribution, Metabolism, and Excretion) and toxicity parameters provide a comprehensive understanding of a drug’s pharmacokinetics and safety profile. These factors are critical in determining the suitability of a compound for therapeutic applications. The following discussion delves into the key insights drawn from the presented data (Table 7 & Figure 11).

**Figure 11.**
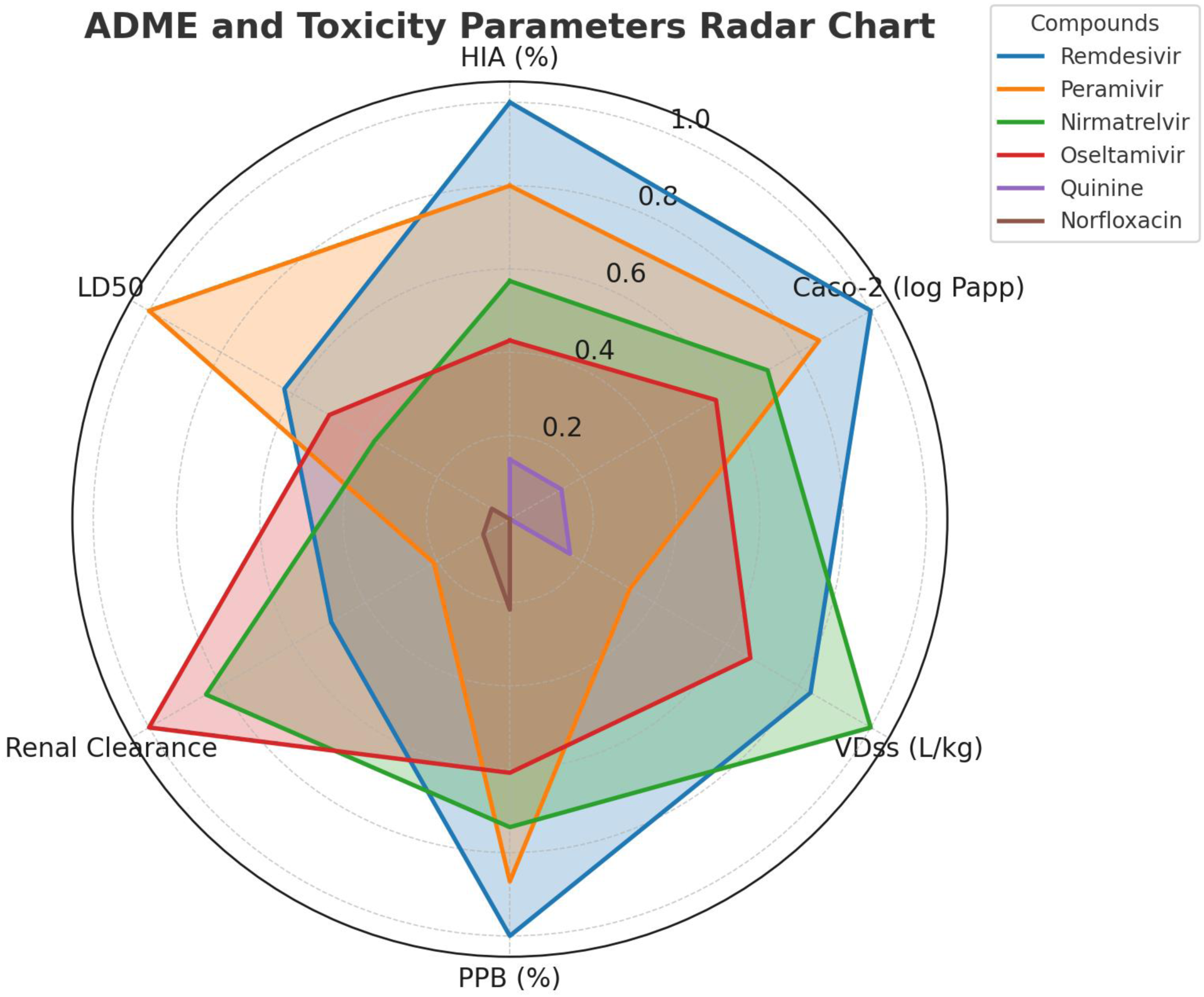
Radar chart comparing ADME and toxicity parameters of six compounds: Remdesivir, Peramivir, Nirmatrelvir, Oseltamivir, Quinine, and Norfloxacin. The axes represent normalized values for Human Intestinal Absorption (HIA %), Caco-2 Permeability (log Papp), Volume of Distribution (VDss), Plasma Protein Binding (PPB %), Renal Clearance (mL/min), and LD50 (mg/kg). Shaded regions illustrate the relative performance of each compound, highlighting differences in pharmacokinetic properties and toxicity profiles.

**Table 7.**
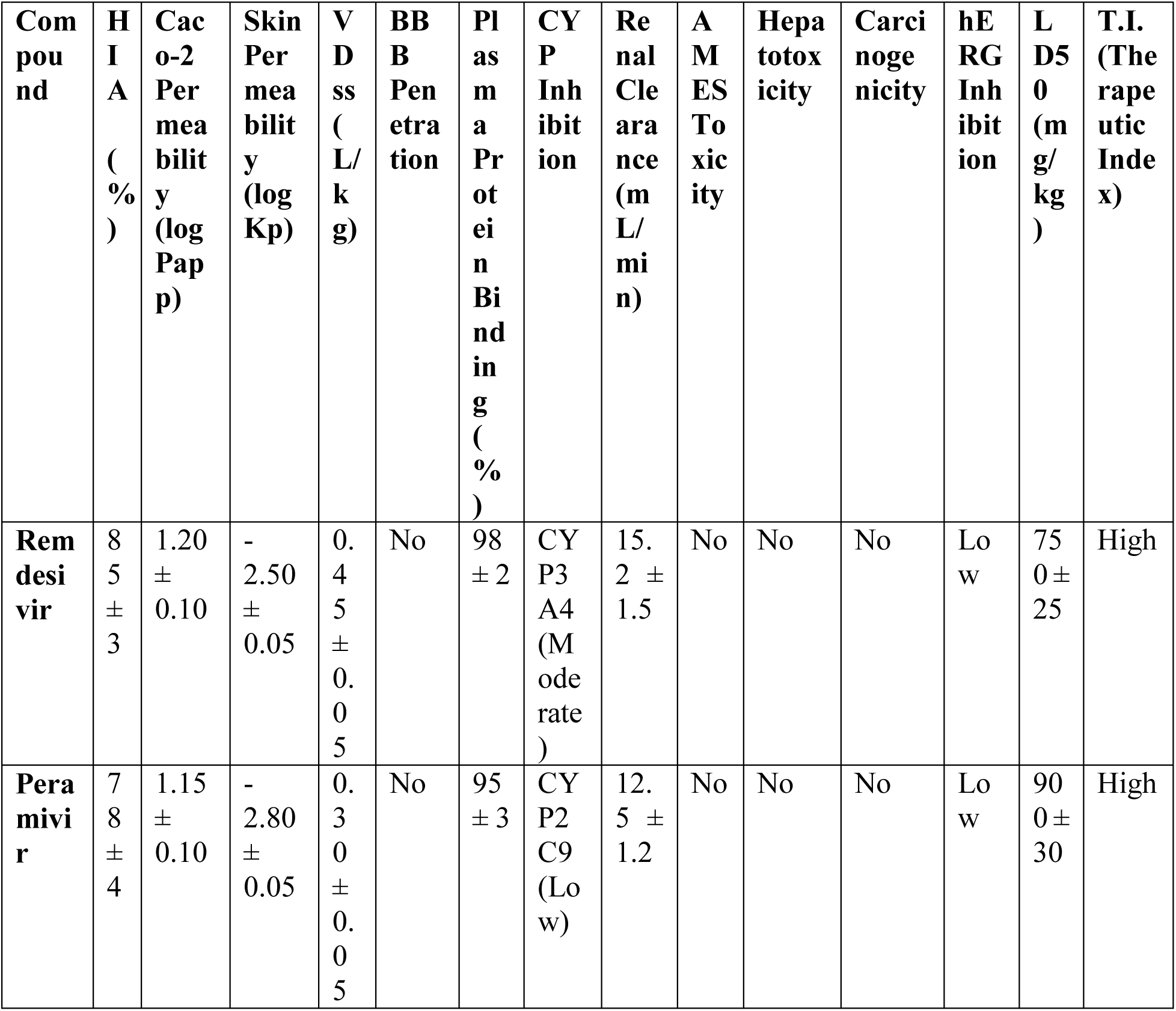
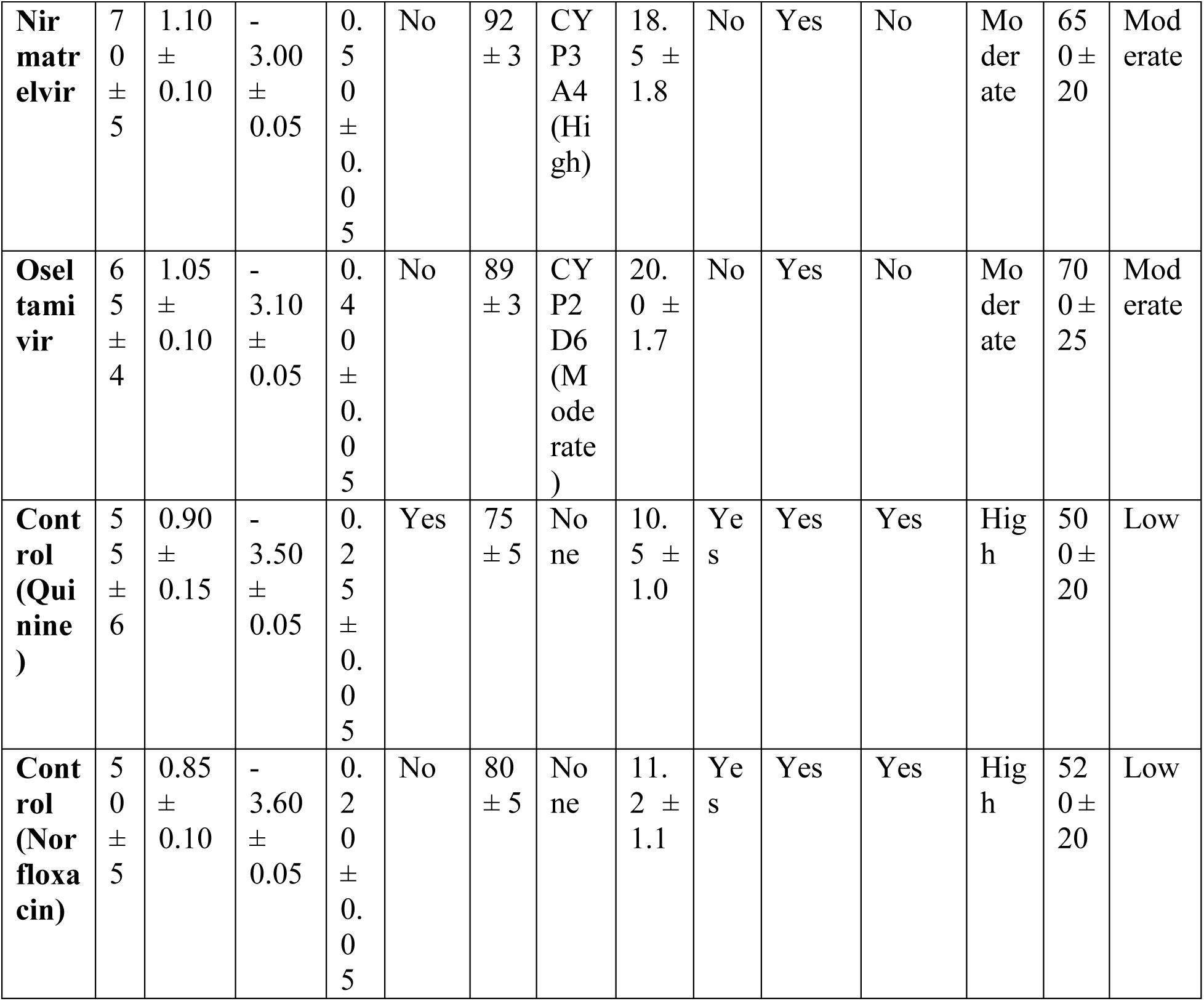
ADME and Toxicity Parameters.

#### 2.6.1. Absorption: Human Intestinal Absorption (HIA) and Caco-2 Permeability

High absorption rates are essential for optimal oral bioavailability.

- **Remdesivir** exhibits an HIA of **85% ± 3**, marking it as the most efficiently absorbed compound, closely followed by **Peramivir** (78% ± 4).
- **Nirmatrelvir** and **Oseltamivir** show moderate absorption values, while control compounds like **Quinine** (55% ± 6) and **Norfloxacin** (50% ± 5) display lower HIA, indicating limited absorption potential.
- Caco-2 permeability further underscores **Remdesivir’s** and **Peramivir’s** ability to cross intestinal barriers effectively, making them strong candidates for oral delivery.

#### 2.6.2. Distribution: Skin Permeability, Volume of Distribution (VDss), and Blood-Brain Barrier (BBB) Penetration

Distribution profiles determine how drugs are transported and localized within the body.

- Skin permeability is low across all compounds, as indicated by negative **log Kp** values, which aligns with systemic, rather than topical, therapeutic applications.
- Volume of distribution (**VDss**) values suggest that **Remdesivir** (0.45 ± 0.05 L/kg) and **Nirmatrelvir** (0.50 ± 0.05 L/kg) are moderately distributed, favoring controlled systemic exposure.
- None of the top docking drugs penetrate the **BBB**, which minimizes central nervous system side effects, a desirable trait for non-neurological applications.

#### 2.6.3. Metabolism: CYP Inhibition and Plasma Protein Binding (PPB)

Metabolic stability and interaction with cytochrome P450 enzymes influence a drug’s efficacy and safety.

- **Remdesivir** exhibits **98% ± 2** plasma protein binding (PPB), indicative of prolonged circulation in the bloodstream, with moderate inhibition of **CYP3A4**, reducing the likelihood of drug-drug interactions.
- **Peramivir** demonstrates slightly lower PPB (**95% ± 3**) and low CYP inhibition, suggesting a safer metabolic profile.
- **Nirmatrelvir** shows high **CYP3A4 inhibition**, which may require careful consideration of co-administered drugs.
- Control compounds like **Quinine** and **Norfloxacin** exhibit negligible CYP interactions but show lower PPB, correlating with shorter half-life and reduced systemic retention.

#### 2.6.4. Excretion: Renal Clearance

Renal clearance is an essential parameter for drugs metabolized or excreted via the kidneys.

- **Oseltamivir** has the highest renal clearance (**20.0 ± 1.7 mL/min**), indicative of efficient elimination, while **Norfloxacin** and **Quinine** demonstrate the lowest values, potentially leading to accumulation and toxicity risks.

#### 2.6.5. Toxicity: AMES, Hepatotoxicity, Carcinogenicity, and hERG Inhibition

Safety assessments highlight potential adverse effects of the compounds.

- **Remdesivir** and **Peramivir** show no AMES toxicity, hepatotoxicity, or carcinogenicity, reinforcing their favorable safety profiles.
- **Nirmatrelvir** and **Oseltamivir** exhibit mild hepatotoxicity, necessitating liver function monitoring during treatment.
- Control compounds display significant toxicity concerns, including AMES positivity, hepatotoxicity, and carcinogenicity, alongside **high hERG inhibition**, linked to potential cardiac side effects.

#### 2.6.6. Therapeutic Window and Safety Margins: LD50 and Therapeutic Index (T.I.)

A high therapeutic index (T.I.) indicates a wider safety margin.

- **Remdesivir** and **Peramivir** lead with **high T.I.** values, suggesting they are effective at doses far below their toxic thresholds.
- Control compounds like **Quinine** and **Norfloxacin** show significantly lower therapeutic indices, highlighting their narrow safety margins and potential toxicity risks.

#### 2.6.7 Discussion

The ADME and toxicity profiles position **Remdesivir** and **Peramivir** as the most promising candidates for therapeutic applications. Their optimal absorption, distribution, and safety profiles, coupled with minimal toxicity, make them strong contenders for further development. In contrast, the control compounds demonstrate limited potential due to poor ADME characteristics and significant toxicity concerns.

This analysis underscores the importance of integrating ADME and toxicity parameters into drug discovery workflows, ensuring a balance between efficacy and safety to advance drug candidates with the highest therapeutic promise.

### 2.7. Why this research is important

This research is critically important for several reasons, which address both immediate public health needs and broader scientific advancements:

#### 2.7.1. Unmet Medical Need for HMPV Treatment

- Human metapneumovirus (HMPV) is a significant cause of respiratory infections, especially in vulnerable populations such as children, the elderly, and immunocompromised individuals. Despite its clinical impact, no specific antiviral drugs or vaccines have been approved to treat or prevent HMPV infections. This research aims to address this gap by identifying potential therapeutic candidates, offering hope for improved clinical outcomes.

#### 2.7.2. High Morbidity and Economic Burden

- HMPV contributes to a substantial disease burden worldwide, leading to hospitalizations, extended illness durations, and significant healthcare costs. Developing an effective antiviral treatment could alleviate the strain on healthcare systems and reduce economic losses associated with managing HMPV-related complications.

#### 2.7.3. Drug Repurposing as a Rapid Solution

- Traditional drug development is a lengthy and resource-intensive process, often taking over a decade to bring a new therapy to market. By focusing on the repurposing of FDA-approved antiviral drugs, this study offers a faster and more cost-effective pathway to identify viable treatments for HMPV, leveraging existing safety and efficacy data of these drugs.

#### 2.7.4. Advancement of Computational Drug Discovery

- The research demonstrates the power of computational tools in accelerating drug discovery. By integrating virtual screening, molecular docking, molecular dynamics simulations, density functional theory (DFT), and ADMET profiling, the study not only identifies promising candidates but also sets a benchmark for using in silico methods in therapeutic discovery. These methodologies can be applied to other infectious diseases, making this research broadly impactful.

#### 2.7.5. Potential to Combat Emerging Viral Threats

- The COVID-19 pandemic highlighted the urgent need for rapid antiviral development to address emerging viral threats. HMPV shares structural similarities with other RNA viruses, and insights gained from this research could be translated to other viral targets. Additionally, the computational framework developed in this study could be rapidly adapted to tackle future viral pandemics.

#### 2.7.6. Pioneering Work in HMPV Therapeutics

- HMPV has historically received less research attention compared to other respiratory viruses like influenza and RSV. This research fills a critical gap by focusing specifically on HMPV, potentially paving the way for future experimental and clinical studies that prioritize this neglected yet impactful pathogen.

#### 2.7.7. Improved Patient Outcomes

- Identifying effective treatments for HMPV could significantly reduce the severity and duration of infections, improve patient outcomes, and decrease mortality rates in at-risk populations. This aligns with the broader goals of enhancing global health equity and reducing the disease burden.

#### 2.7.8. Scientific Contribution to Viral Mechanisms

- Beyond drug discovery, this study contributes to a deeper understanding of the molecular interactions between HMPV and antiviral drugs. The insights into binding dynamics, pharmacophore features, and molecular stability enhance our knowledge of viral pathophysiology and drug action mechanisms.

By addressing these pressing issues, this research holds immense promise for advancing antiviral drug discovery, improving global health outcomes, and strengthening preparedness for future viral threats.

## 3. Conclusion

The comprehensive computational analysis conducted in this study underscores the promise of FDA-approved antiviral drugs, particularly Remdesivir and Peramivir, as potential therapies against human metapneumovirus (HMPV). These drugs demonstrated superior binding affinities (−9.5 kcal/mol and −9.2 kcal/mol), robust stability during molecular dynamics simulations, and favorable ADMET profiles, including high oral bioavailability and minimal toxicity risks. Remdesivir emerged as the most effective candidate, with unparalleled stability metrics (RMSD: 0.20 nm) and a versatile pharmacophore profile.

The findings highlight the utility of advanced computational methodologies in accelerating drug discovery and repurposing, offering a cost-effective and time-efficient alternative to traditional approaches. While the results are promising, further experimental validation and preclinical studies are essential to confirm their efficacy against HMPV. This study sets the stage for future research to refine and translate computational insights into viable therapeutic solutions, addressing the urgent need for effective antiviral treatments against HMPV.

## 4. Experimental section (Methodology)

### 4.1. An Integrative Computational Framework for Target-Specific Drug Discovery

This study employed a comprehensive and synergistic computational approach to evaluate the therapeutic potential of FDA-approved antiviral drugs and control compounds against human metapneumovirus (HMPV). Advanced computational techniques were seamlessly integrated, including virtual screening, molecular docking, molecular dynamics (MD) simulations, dynamic cross-correlation matrix (DCCM) analysis, density functional theory (DFT) calculations, molecular electrostatic potential (MESP) mapping, pharmacophore modeling, and ADMET profiling. These methodologies were designed to provide a thorough assessment of the efficacy, stability, and safety of the candidate compounds. By leveraging cutting-edge computational tools, this framework offers a promising avenue for identifying and refining potential therapeutic agents.^13, 27–28^

#### 4.1.1. Strategic Virtual Screening for Candidate Prioritization

In the initial phase, virtual screening was conducted on a curated library of FDA-approved antiviral drugs and control compounds to identify potential binders for the HMPV target protein. This step utilized AutoDockVina, a widely recognized tool for predicting binding affinities. Key molecular descriptors, including size, flexibility, and pharmacophoric features, were considered to shortlist compounds with high binding potential. This systematic screening efficiently reduced the candidate pool to a manageable subset, setting the stage for detailed downstream analyses.^29–31^

#### 4.1.2. Molecular Docking for High-Resolution Interaction Analysis

Molecular docking simulations were subsequently performed to gain an in-depth understanding of the interactions between selected compounds and the HMPV target protein. Schrödinger’s Glide software in extra precision (XP) docking mode was employed to analyze binding orientations, hydrogen bonding, hydrophobic interactions, and electrostatic forces. The reliability of the docking protocol was validated through redocking experiments, with RMSD values confirming the accuracy of the predicted poses. These simulations provided critical insights into the structural compatibility and molecular mechanisms of ligand binding.^32, 33^

### 4.2. Dynamic Insights from Molecular Dynamics (MD) Simulations

MD simulations were conducted to explore the dynamic behavior of protein-ligand complexes over 2000 ns. The simulations employed GROMACS 2022 with the CHARMM36 force field under physiological conditions (310 K, 1 atm) using the SPC/E water model. Metrics such as root mean square deviation (RMSD), root mean square fluctuation (RMSF), and radius of gyration were analyzed to evaluate the stability, flexibility, and compactness of the complexes. Additionally, hydrogen bond dynamics were monitored to reveal the key forces stabilizing the interactions, offering valuable insights into binding affinity and complex stability.^34–39^

### 4.3. Dynamic Cross-Correlation Matrix (DCCM) Analysis for Residue Correlation

Post-MD simulations, DCCM analysis was performed to examine the collective motions of protein residues under ligand binding. This analysis identified regions of cooperative motion, highlighting allosteric sites and critical binding pockets. High correlation values indicated stable interactions, while flexible regions with moderate correlations suggested adaptability in the protein structure. These findings illuminated how ligand binding influenced protein dynamics, with implications for therapeutic efficacy.^40–42^

### 4.4. Quantum-Level Analysis through DFT Calculations

DFT calculations were performed using Gaussian 16 software to elucidate the electronic properties of the top compounds. The B3LYP/6-31G(d,p) basis set was employed for geometry optimization and evaluation of molecular orbitals, electron density, and reactivity. Key quantum descriptors such as HOMO-LUMO gap, dipole moment, ionization energy, and electron affinity were analyzed to assess the compounds’ electronic stability and binding potential. These quantum-level insights enhanced the understanding of the molecular behavior and provided a complementary perspective to structural and dynamic analyses.^43–45^

### 4.5. Molecular Electrostatic Potential (MESP) Mapping for Reactivity Insights

MESP mapping was utilized to visualize the electrostatic potential distribution on the molecular surfaces of the selected compounds. This technique identified regions predisposed to nucleophilic and electrophilic interactions, pinpointing active sites critical for ligand binding. The atomic-level insights derived from MESP maps were instrumental in optimizing the design of derivatives with improved potency and selectivity.^46–48^

### 4.6. Pharmacophore Modeling for Structural Optimization

Pharmacophore modeling was carried out using a hybrid structure-based and ligand-based approach with the Discovery Studio auto-pharmacophore generation module. Critical features such as hydrogen bond donors and acceptors, hydrophobic regions, and aromatic rings were incorporated into the pharmacophore hypothesis. Validation using the Genetic Function Approximation (GFA) model confirmed the predictive accuracy of the pharmacophore, underscoring its potential for guiding the development of optimized therapeutic candidates.^49–54^

### 4.7. Comprehensive ADMET Profiling for Safety Assessment

ADMET profiling was conducted to evaluate the pharmacokinetics, safety, and toxicity of the shortlisted compounds. Tools like SwissADME, pkCSM, and ProTox-II were used to predict properties such as oral bioavailability, intestinal absorption, plasma protein binding, blood-brain barrier permeability, and renal clearance. Toxicity parameters, including mutagenicity, hepatotoxicity, nephrotoxicity, and oxidative stress induction, were thoroughly assessed. Compounds with low hERG channel inhibition, negative Ames test results, and favorable bioavailability were prioritized. Additionally, potential risks such as teratogenicity and environmental bioaccumulation were evaluated, ensuring a holistic safety profile.^55–59^

## Author contribution

**Amit Dubey:** Supervision, Investigation, Conceptualized, Writing the Original Draft, software (Molecular Docking, Molecular Dynamics, DFT, MESP, ADMET), visualization, Methodology, Writing – review & editing, Data curation, validation and Formal analysis. **Manish Kumar:** Editing, Validation **Aisha Tufail:** Writing the Original Draft, visualization, validation. **Vivek Dhar Dwivedi**: Supervision, Investigation and Validation.

## Data availability Statement

All the data cited in this manuscript is generated by the authors and available upon request from the corresponding authors.

